# Differential selection between sexes and the evolution of recombination in haplodiploids

**DOI:** 10.64898/2026.06.29.735359

**Authors:** Vinayak Patel, Denis Roze

**Affiliations:** Indian Institute of Science Education and Research (IISER), Pune 411008, India; Sorbonne Université, CNRS, UMR 7144 AD2M, DiSEEM, Station Biologique de Roscoff, 29680 Roscoff, France

**Keywords:** epistasis, eusociality, meiosis, modifier model, multilocus model

## Abstract

Eusocial Hymenoptera present the highest known recombination rates among metazoans, which evolved several times independently among bees, ants and wasps. Several hypotheses have been proposed to explain this observation, including stronger selection for recombination caused by coevolving parasites and pathogens, and strong sexual selection among haploid males due to male-biased sex ratios among reproductive individuals. In this article, we explore the effects of haplodiploidy and differential selection between sexes on the evolution of recombination, by analyzing a three-locus model in which selection for recombination stems from negative epistasis between selected loci. Our analytical predictions are compared with the results of individual-based simulations in which deleterious mutations occur along a linear chromosome. Our results show that, at mutation-selection balance for deleterious alleles, increasing the strength of selection against deleterious alleles (due to the effect of male haploidy and/or sexual selection) tends to reduce selection for recombination. However, an increase in the overall magnitude of negative epistasis (which may also be due to male haploidy and/or sexual selection) combined with the fact that recombination only occurs in females may increase selection for recombination substantially. Our model also shows that, in conditions favoring recombination, increasing recombination in meioses leading to parthenogenetic ovules (and male offspring) may yield stronger benefits than in meioses leading to fertilized ovules (and female offspring).

## INTRODUCTION

While genetic recombination represents a fundamental aspect of sexual reproduction, generating new genotypic combinations by crossing-over among homologous chromosomes, recombination rates (measured in cM/Mb) vary over several orders of magnitude among eukaryotes (Stapley et al., 2017). This variation is largely due to differences in the physical size of chromosomes, the number of crossovers per chromosome staying comprised between 1 and 3 or 4 in the majority of species (Fernandes et al., 2018; Brazier and Glémin, 2022). Yet, heritable genetic variation for recombination rates has been demonstrated in an increasing number of species (Johnston, 2024; Payseur, 2025) and recombination can evolve over relatively short timescales, leading to differences in recombination landscapes between closely related species, or between populations from the same species (e.g., Brand et al., 2018; Samuk et al., 2020). A change in recombination rate may have direct fitness effects (for example, by affecting the segregation of homologous chromosomes during meiosis I, or the probability of ectopic recombination between repeated sequences), and may also be indirectly selected, through its effect on genetic associations between selected loci. In particular, higher recombination increases the variance in fitness among offspring in the presence of negative linkage disequilibrium (LD) between selected loci, thereby enhancing the efficiency of natural selection. In that case, a mutation increasing recombination may rise in frequency by hitchhiking with the beneficial genotypes it has contributed to create (Otto and Lenormand, 2002; Agrawal, 2006). Negative LD may result from negative epistasis between selected loci (Charlesworth, 1990; Barton, 1995), or from the Hill-Robertson effect in finite populations (Hill and Robertson, 1966; Felsenstein, 1974; Otto and Barton, 1997; Barton and Otto, 2005; Keightley and Otto, 2006; Roze, 2021). Additionally, recombination may increase the average fitness of offspring under some forms of spatially or temporally varying selection (Barton, 1995; Lenormand and Otto, 2000; Gandon and Otto, 2007; Salathé et al., 2009).

Assessing the evolutionary causes of changes in recombination rates within natural populations — and the relative importance of direct and indirect effects — is difficult. Given that linkage maps are becoming available for a rapidly increasing number of species, valuable insights may be gained by comparative analyses, testing for possible associations between the ecology, life cycle or reproductive system of organisms with their recombination rates, in ways that align with theoretical predictions. For example, self-fertilizing species of Angiosperms tend to display higher recombination rates than outcrossing species (Roze and Lenormand, 2005; Ross-Ibarra, 2007; Brazier et al., 2025), which may be explained by the fact that self-fertilization reduces effective recombination rates (since recombination has no effect in homozygous individuals), increasing the magnitude of genetic associations and selecting more strongly for recombination in the presence of negative LD generated by negative epistasis or by the Hill-Robertson effect (Stetsenko and Roze, 2022).

Interestingly, eusocial Hymenoptera display very high rates of recombination, associated with three independent origins of eusociality in bees, ants and wasps (DeLory et al., 2024). In particular, honey bees exhibit the highest known rates of recombination in metazoans, with 5.7 crossovers per chromosome and 20.8 cM/Mb on average (Beye et al., 2006; Wilfert et al., 2007; Rueppell et al., 2016; Wallberg et al., 2015), while eusocial wasps, ants, bumble bees and stingless bees also show high recombination rates (e.g., Sirviö et al., 2006, 2010, 2011; Kawakami et al., 2019; Waiker et al., 2021). Hymenoptera have a haplodiploid life cycle: females are diploid, while males are haploid and produced by the parthenogenetic development of unfertilized ovules. Several authors proposed the idea that higher recombination rates may have been favored to compensate for the fact that recombination only occurs in females (e.g., Wilfert et al., 2007), but this does not explain the high recombination rates observed in eusocial species, since solitary bees and wasps (which are also haplodiploid) do not exhibit particularly high recombination rates (Niehuis et al., 2010; Jones et al., 2019; DeLory et al., 2024).

Several classes of hypotheses have been proposed to explain the evolution of high recombination rates in eusocial species (DeLory et al., 2024). A first is based on the idea that increasing genotypic diversity within colonies by recombination provides an advantage at the colony level, either by allowing the colony to better resist against parasites and pathogens, or by increasing the efficiency of division of labor (Gadau et al., 2000; Schmid-Hempel, 2000). However, another means of increasing within-colony diversity is polyandry, which occurs frequently in eusocial species (Schmid-Hempel and Crozier, 1999; Schmid-Hempel, 2000; Rueppell et al., 2012). A second hypothesis proposes that increased recombination helps to prevent nepotistic conflicts within colonies, by reducing the variance in relatedness (averaged over the whole genome) between individuals from the same colony (Sherman, 1979). Templeton (1979) presented an alternative argument based on the idea that reducing the variance in relatedness within colonies decreases the variance of inclusive fitness benefits of social traits (when these traits are polygenic), which should be favored by selection for reduced variance in fitness (Gillespie, 1974). Finally, a third class of hypotheses proposes that the benefits of recombination in terms of increased efficiency of natural selection are particularly strong in eusocial species, either because (i) selective pressures generated by coevolving parasites are high, due to the fact that parasite transmission is facilitated by colonial life (Seger and Hamilton, 1988; Sirviö et al., 2006), (ii) the evolution of caste specialization favored increased recombination between genes involved in caste phenotypes (Kent et al., 2012; Kent and Zayed, 2013), or because (iii) eusocial species have particularly low effective population sizes (Romiguier et al., 2014; Weyna and Romiguier, 2021), increasing the strength of selection for recombination generated by the Hill-Robertson effect.

These hypotheses predict that higher recombination rates should evolve in eusocial species, including species in which males are diploid. However, a recent study by Everitt et al. (2025) showed that termites (which are eusocial, but in which both females and males are diploid) do not display high recombination rates. Based on these results, Everitt et al. (2025) proposed an alternative hypothesis to explain the high recombination rates of eusocial Hymenoptera, involving the effect of sexual selection among males. According to this hypothesis, sexual selection is particularly intense in these species due to strong male-biased sex ratios among reproductive offspring. This strong sexual selection favors high recombination rates in order to increase the variance in fitness among male offspring, thereby increasing the chance to produce a high-fitness genotype — this implicitly assumes a source of negative LD between selected loci (either negative epistasis or the Hill-Robertson effect), so that recombination increases the variance in fitness. By contrast, the sex ratio is relatively even in nonsocial Hymenoptera and in termites, and sexual selection may therefore be less strong in these species.

Using current models on the evolution of recombination to assess the combined effects of female-limited recombination, male haploidy and stronger selection among males on selection for recombination is difficult, as these models do not consider the case of haplodiploid life cycles. In this article, we explore how haplodiploidy and differential selection between the sexes affect the evolution of recombination. We develop and analyze deterministic models in which selection for recombination results from negative epistasis between deleterious mutations, and compare our analytical predictions to the results of individual-based, multilocus simulations. In order to compare the strength of selection for recombination in diploids and haplodiploids, we use Lenormand’s (2003) model to obtain predictions for the diploid case when selection and recombination may differ between the sexes (this model is extended to obtain results that can be extrapolated to the case where deleterious mutations occur along an entire chromosome). Our results show that female-limited recombination increases the overall strength of indirect selection on recombination in haplodiploids, by increasing the magnitude of genetic associations between loci. Increasing the overall strength of selection against deleterious alleles in males (either due to male haploidy or sexual selection) generally reduces the strength of selection for recombination. However, stronger negative epistasis in males (which may again be due to male haploidy or sexual selection) increases selection for recombination.

## METHODS

### Genetic architecture and life cycle

Our three-locus model considers two selected loci at which deleterious alleles *a* and *b* occur by mutation from wild-type alleles *A* and *B* at a rate *u* per generation, and a recombination modifier locus with two alleles *M* and *m*. All loci are assumed to be present on autosomes. In order to simplify the notation, the letters *m, a* and *b* will also be used to identify the three loci. All results given in the article are valid for any ordering of the three loci along the chromosome (*m* − *a* − *b* or *a* − *m* − *b*). The parameters and variables of the model are summarized in Tables 1 and 2.

**Table 1.** Parameters of the model.

|  |  |
| --- | --- |
| $s^{\varphi}, s^{\sigma}$ | Selection coefficients of deleterious alleles in females and males |
| $h^{\varphi}, h^{\sigma}$ | Dominance coefficients of deleterious alleles in females and males (when diploid) |
| $e_{\text{axa}}^{\varphi}, e_{\text{axa}}^{\sigma}$ | Epistasis between heterozygous deleterious mutations in diploid females and males |
| $e^{\sigma}$ | Epistasis between deleterious mutations in haploid males |
| $r_{ij}^{\varphi}, r_{ij}^{\sigma}$ | Recombination rate between loci $i$ and $j$ in females and males (diploid model) |
| $\delta r_{ij}^{\varphi}, \delta r_{ij}^{\sigma}$ | Modifier effect on the recombination rate between loci $i$ and $j$ in females and males (diploid model) |
| $r_{ij}^{\varphi\varphi}, r_{ij}^{\varphi\sigma}$ | Recombination rate between loci $i$ and $j$ in meioses giving rise to fertilized and unfertilized ovules (haplodiploid model) |
| $\delta r_{ij}^{\varphi\varphi}, \delta r_{ij}^{\varphi\sigma}$ | Modifier effect on the recombination rate between loci $i$ and $j$ in meioses giving rise to fertilized and unfertilized ovules (haplodiploid model) |
| $h_m$ | Dominance coefficient of modifier allele $m$ |
| $R^{\varphi}, R^{\sigma}$ | Chromosome map length in females and males (diploid model) |
| $\delta R^{\varphi}, \delta R^{\sigma}$ | Modifier effect on chromosome map length in females and males (diploid model) |
| $R^{\varphi\varphi}, R^{\varphi\sigma}$ | Chromosome map length in meioses giving rise to fertilized and unfertilized ovules (haplodiploid model) |
| $\delta R^{\varphi\varphi}, \delta R^{\varphi\sigma}$ | Modifier effect on chromosome map length in meioses giving rise to fertilized and unfertilized ovules (haplodiploid model) |
| $u$ | Deleterious mutation rate (per locus) |
| $U$ | Deleterious mutation rate (per chromosome) |
| $c$ | Direct fitness cost of recombination (per crossover) |
| $N$ | Population size (in the simulations) |

**Table 2.** Model variables.

|  |  |
| --- | --- |
| $p_{i,\emptyset}^{\circ}, p_{\emptyset,i}^{\circ}$ | Frequency of allele $i$ on the maternally and paternally inherited chromosomes of diploid females |
| $p_{i,\emptyset}^{\sigma}, p_{\emptyset,i}^{\sigma}$ | Frequency of allele $i$ on the maternally and paternally inherited chromosomes of diploid males |
| $p_i^{\sigma}$ | Frequency of allele $i$ in haploid males |
| $p_i = \frac{1}{2} (p_{i,\emptyset}^{\circ} + p_{\emptyset,i}^{\circ})$ | Average frequency of allele $i$ in the diploid model (same in females and males at the start of a generation) |
| $\tilde{p}_i = \frac{1}{3} (p_{i,\emptyset}^{\circ} + p_{\emptyset,i}^{\circ} + p_i^{\sigma})$ | Weighted frequency of allele $i$ in the haplodiploid model |
| $\nabla_i = p_{i,\emptyset}^{\circ} - p_{\emptyset,i}^{\circ}$ | Difference in frequency of allele $i$ on maternally and paternally inherited chromosomes |
| $D_{\mathbb{U},\mathbb{V}}^{\circ}, D_{\mathbb{U},\mathbb{V}}^{\sigma}$ | Genetic association between sets of alleles $\mathbb{U}$ and $\mathbb{V}$ present on the maternally and paternally inherited genomes of diploid females and males |
| $D_{\mathbb{U}}^{\sigma}$ | Genetic association between the set of alleles $\mathbb{U}$ in haploid males |

Females are diploid and produced by fertilization while males are haploid and result from the parthenogenetic development of unfertilized ovules. The fitness of females as a function of their genotype at the selected loci is given in Table 3 (which takes the same form as in Roze and Lenormand, 2005 and Stetsenko and Roze, 2022): *s*^♀^ and *h*^♀^ are the selection and dominance coefficients of deleterious alleles (*a* and *b*) in females, while epistasis between deleterious alleles is decomposed into three terms corresponding to the effect of the interaction between two (additive-by-additive epistasis, 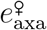), three (additive-by-dominance epistasis, 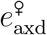) and four (dominance-by-dominance epistasis, 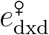) deleterious alleles at the two selected loci. The fitnesses of *AB, Ab, aB* and *ab* males are given by 1, 1 − *s*^♂^, 1 − *s*^♂^ and (1 − *s*^♂^)^2^ + *e*^♂^, respectively (*e*^♂^ thus measures the effect of epistasis between deleterious alleles in haploid males). We assume random mating, *h*^♀^ significantly greater than zero and *u « s*^♀^, *s*^♂^, so that the frequencies of deleterious alleles remain small, *a* and *b* being mostly present in the heterozygous state in females: therefore, selection among females mainly depends on *s*^♀^*h*^♀^ and 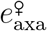. The number of gametes produced by each individual is proportional to its fitness, meiosis and recombination only occurring in females. Because recombination rates in Hymenoptera are typically measured in sons, it is unknown whether recombination rates are similar in meioses leading to daughters and in those leading to sons (see Discussion). Therefore, for generality we assume that recombination rates may differ between meioses giving rise to ovules that will be fertilized (generating female offspring) and meioses giving rise to ovules that will develop parthenogenetically (generating male offspring). The recombination rate between loci *i* and *j* in *MM*, *Mm* and *mm* females is denoted 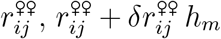 and 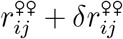 (respectively) in the first type of meiosis (giving rise to female offspring) and 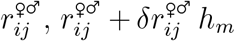 and 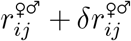 in the second type (giving rise to male offspring). Because recombination between the modifier and each selected locus only has an effect on the gametes produced by *Mm* heterozygous females, 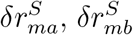 (where *S* may be ♀♀ or ♀♂) do not generate indirect selection at the modifier locus, which is entirely driven by 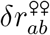 and 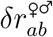 (e.g., Barton, 1995; Otto and Barton, 1997). Male and female gametes fuse at random to produce the females of the next generation, while males are produced from unfertilized ovules.

**Table 3.** Fitness matrix for diploid females. Under random mating and when deleterious alleles *a* and *b* stay rare in the population, the effect of selection is determined mostly by *s*^♀^*h*^♀^ and 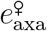. In the diploid model, the fitnesses of males are given by the same expressions, replacing female by male symbols. In the haplodiploid model, the fitnesses of *AB, Ab, aB* and *ab* males are given by 1, 1 − *s*^♂^, 1 − *s*^♂^ and (1 − *s*^♂^)^2^ + *e*^♂^, respectively.

| | $AA$ | $Aa$ | $aa$ |
| --- | --- | --- | --- |
| $BB$ | 1 | $1 - s^{\varphi}h^{\varphi}$ | $1 - s^{\varphi}$ |
| $Bb$ | $1 - s^{\varphi}h^{\varphi}$ | $(1 - s^{\varphi}h^{\varphi})^2 + e_{\text{axa}}^{\varphi}$ | $(1 - s^{\varphi}h^{\varphi})(1 - s^{\varphi}) + 2e_{\text{axa}}^{\varphi} + e_{\text{axd}}^{\varphi}$ |
| $bb$ | $1 - s^{\varphi}$ | $(1 - s^{\varphi}h^{\varphi})(1 - s^{\varphi}) + 2e_{\text{axa}}^{\varphi} + e_{\text{axd}}^{\varphi}$ | $(1 - s^{\varphi})^2 + 4e_{\text{axa}}^{\varphi} + 4e_{\text{axd}}^{\varphi} + e_{\text{dxd}}^{\varphi}$ |

In order to compare selection for recombination in haplodiploids and diploids, we also analyzed a diploid model (similar to the model analyzed by Lenormand, 2003), in which selection and recombination may differ between the sexes. In this case, the fitness of males is also given by Table 3, replacing female by male symbols. We again assume that deleterious alleles stay rare (*h*^♀^, *h*^♂^ significantly greater than zero, *u* ≪ *s*^♀^, *s*^♂^), so that selection mainly depends on *s*^♀^*h*^♀^, *s*^♂^*h*^♂^, 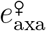 and 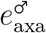. During meiosis, the recombination rate between loci *i* and *j* is given by 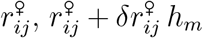 and 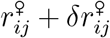 in *MM*, *Mm* and *mm* females (respectively), and by 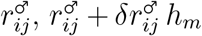 and 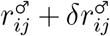 in *MM*, *Mm* and *mm* males. Gametes fuse at random, and each new individual becomes female or male with probability 0.5, independently of its genotype at loci *m, a* and *b*. The analysis of the diploid model is presented in Supplementary File S1.

### Allele frequencies and genetic associations

Following Barton and Turelli (1991) and Kirkpatrick et al. (2002), we define indicator variables *X*_*i*,∅_ and *X*_∅,*i*_ that equal 1 if allele *i* is present in the maternally (for *X*_*i*,∅_) or paternally (*X*_∅,*i*_) inherited genome of a diploid individual, and 0 otherwise (*i* may be *m, a*, or *b*). The frequency of allele *i* in the maternally inherited genomes of females is thus given by 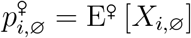, where E^♀^ stands for the average over all females, while the frequency of *i* in the paternally inherited genome of females is 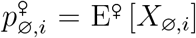. We then define an indicator variable *X*_*i*_ that equals 1 if allele *i* is present in a given haploid male, and 0 otherwise, the frequency of allele *i* in males being given by 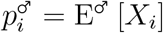.

At the start of a generation, allele frequencies may differ between males and females. Throughout the article, we use weighted allele frequencies, denoted as 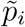, and defined so that the change in 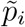 over one generation equals zero in the absence of selection. For this, allele frequencies in females and males must be weighted by the reproductive value of each sex, corresponding to the asymptotic probability that a gene sampled in the present population was in an individual of that sex a long time ago in the past (Taylor, 1990). One can show that the reproductive values of females and males are 2*/*3 and 1*/*3 independently of the sex ratio (e.g., Roze and Rousset, 2004), so that 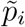 is given by:

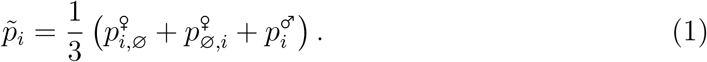

Centered variables measured in females are defined as 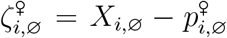 and 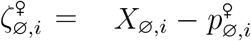, while centered variables measured in haploid males are defined as 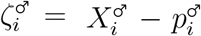. As in Kirkpatrick et al. (2002), the genetic association between sets of alleles U and V present on the maternally and paternally inherited genome of a female is defined as:

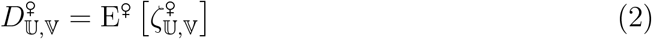

with:

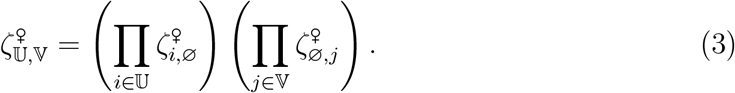

Genetic associations among haploid males are defined as:

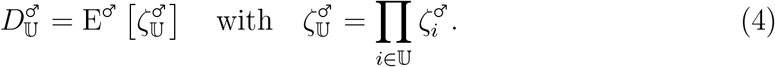

### Selection coefficients and change in frequency at the modifier locus

The method of Kirkpatrick et al. (2002) can be used to compute the effect of selection on allele frequencies and genetic associations. For this, the fitness of a female relative to the average fitness of females in the population is written under the form:

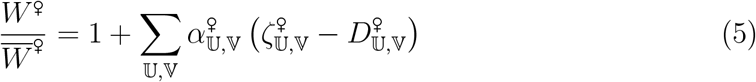

where the sum is over all possible pairs of subsets of alleles U and V at selected loci, each taken in *{*∅, *a, b, ab}*, and where 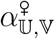 represents the effect of selection acting on the sets of alleles U and V present on the maternally and paternally inherited haplotypes of a female. As the deleterious alleles *a* and *b* are mostly present in the heterozygous state under our assumptions, selection will be mainly driven by the coefficients 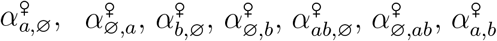 and 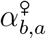. Furthermore, because we assume no sex-of-origin effect at selected loci, we have 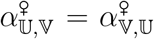, and we define the coefficients 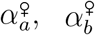 and 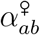 as 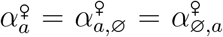 (and similarly for 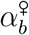) and 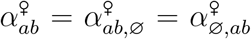. The coefficients 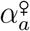 and 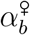 measure the effective strength of directional selection against the deleterious alleles *a* and *b* in females, while 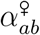 and 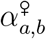 represent epistasis between *a* and *b* present on the same or on different haplotypes (cis and trans epistasis), measured as deviations from additive effects. In our derivations we will assume that selection against deleterious alleles is weak (*s*^♀^ and *s*^♂^ of order *ϵ*, where *ϵ* is a small term), while epistasis terms are weaker (of order *ϵ*^2^). In that case, and given that deleterious alleles stay at low frequency, we have to leading order in *ϵ* (e.g., Barton and Turelli, 1991; Abu Awad and Roze, 2020):

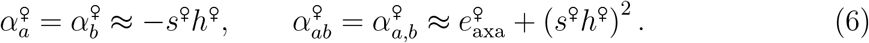

To represent selection among haploid males, 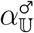 coefficients are defined from:

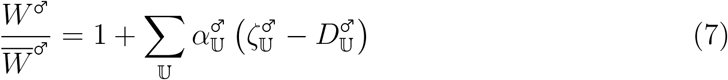

yielding three coefficients 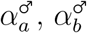 and 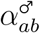. To leading order in *ϵ*, we have:

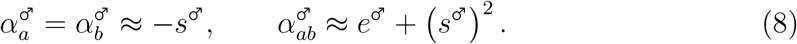

Recurrence equations describing the changes in allele frequencies due to selection can be obtained from equation 10 in Kirkpatrick et al. (2002). In a diploid model, genetic associations are the same in females and males at the start of a generation (as we assume that all loci are autosomal), and one obtains the following expression for the change in frequency of the modifier allele *m* over one generation (Barton, 1995; Lenormand, 2003):

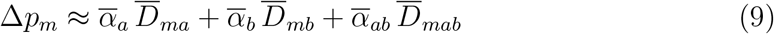

with 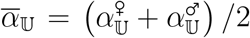 and 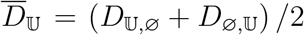. Under negative epistasis, the term 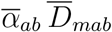 generally disfavors recombination, corresponding to the effect of the decreased average fitness of offspring caused by recombination (recombination load; e.g., Barton, 1995; Charlesworth and Barton, 1996; Otto and Lenormand, 2002), while the term 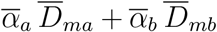 generally favors recombination, corresponding to the effect of increased variance in fitness among offspring (thereby increasing the efficiency of natural selection) due to recombination. Under positive epistasis, all terms disfavor recombination, as recombination decreases both the mean fitness and the variance in fitness among offspring (Barton, 1995).

In haplodiploids, genetic associations may differ between females and males, and one obtains that the change in weighted frequency of allele *m* (defined by equation 1) over one generation is given by:

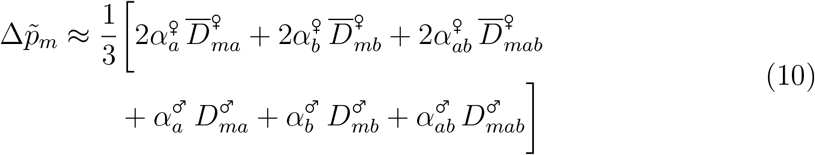

with 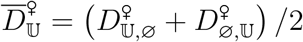. Equation 10 shows that the effects of indirect selection in diploid females and in haploid males are weighted by factors 2*/*3 and 1*/*3, respectively.

### Quasi-linkage equilibrium approximation

The effect of selection on the changes in genetic associations is obtained using equations 9 and 15 in Kirkpatrick et al. (2002), while the method of Barton (1995) can be used to compute the effect of recombination (including the effect of the recombination modifier). As in Barton (1995), we assume that the modifier has a weak effect on recombination rates, and compute all terms to the first order in the modifier effect. When allele frequencies change slowly relative to the rate at which genetic associations decay due to recombination, the quasi-linkage equilibrium (QLE) approximation can be used to compute genetic associations in terms of allele frequencies and of the different parameters of the model (Barton and Turelli, 1991; Nagylaki, 1993; Barton, 1995). In general, this requires that selection is sufficiently weak relative to recombination. However, when deleterious alleles have reached mutation-selection balance, allele frequencies only change due to the modifier effect, and the QLE approximation only requires that the modifier effect is small relative to recombination: therefore, approximations can be obtained for the case where selection coefficients and recombination rates are of the same order of magnitude (e.g., Roze, 2014; Abu Awad and Roze, 2020; Stetsenko and Roze, 2022). Throughout the article, we assume that deleterious alleles stay rare in the population and compute all terms to leading order in the frequencies of *a* and *b*.

### Multilocus extrapolation

As in previous works (Roze, 2021; Stetsenko and Roze, 2022), the result from our three-locus model can be extrapolated to the case where deleterious alleles may occur at a large number of possible loci along a chromosome, at a rate *U* per chromosome per generation. In that case, we assume for simplicity that mutations and crossovers occur at uniform rates along the chromosome (without crossover interference), that all mutations have the same fitness effects and dominance coefficients, and that epistasis is the same between all pairs of mutations. When the mean number of mutations per chromosome is large, approximations for 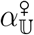 and 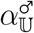 coefficients become:

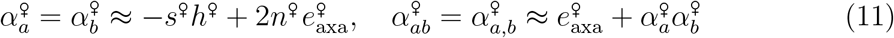

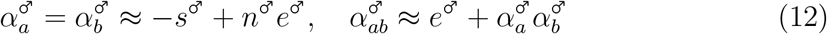

where *n*^♀^ and *n*^♂^ are the mean numbers of mutations per chromosome in females and males, respectively (these numbers are approximately equal when selection is weak).

In this multilocus extrapolation, we assume that the modifier affects the genetic map length of the chromosome (average number of crossovers at meiosis). For this, the map lengths of *MM*, *Mm* and *mm* females are denoted *R*^♀♀^, *R*^♀♀^ + *δR*^♀♀^ *h*_*m*_ and *R*^♀♀^ + *δR*^♀♀^ in meioses leading to daughters, and *R*^♀♂^, *R*^♀♂^ + *δR*^♀♂^ *h*_*m*_ and *R*^♀♂^ + *δR*^♀♂^ in meioses leading to sons. Recombination rates between loci can be computed as a function of their relative positions along the chromosome using Haldane’s mapping function (Haldane, 1919). The result from the three-locus model can then be integrated over all possible positions of deleterious alleles to compute the change in frequency of a recombination modifier allele as a function of its effects on map length in meioses leading to daughters and sons (*δR*^♀♀^, *δR*^♀♂^) and its position along the chromosome (note that this extrapolation neglects the possible effects of interactions between more than two selected loci). A similar extrapolation was performed in the case of the diploid model, assuming that the modifier may have different effects *δR*^♀^ and *δR*^♂^ on the female and male map lengths (*R*^♀^, *R*^♂^).

### Simulations

Individual-based, multilocus simulations were used to check our analytical results in the case of recombination modifiers affecting the map length of a whole chromosome along which deleterious alleles occur at a rate *U* per generation (our simulation programs are written in C++ and available from *Zenodo*, https://doi.org/10.5281/zenodo.20594388). The simulation procedure is described in Supplementary File S2. Simulations were performed for both the haplodiploid and the diploid cases. In both models, chromosome map length is controlled by either one or two modifier loci. In the one-modifier case, map length is the same in males and females (in the diploid model) or in meioses giving rise to daughters and sons (in the haplodiploid model), and the modifier locus is located at the mid-point of the chromosome. In the two-modifier case, one modifier locus controls map length in female meioses (diploid model) or in meioses giving rise to daughters (haplodiploid model), while the other modifier controls map length in male meioses or in meioses giving rise to sons. In that case, the modifier loci are located at 1*/*3 and 2*/*3 of the length of the chromosome. As in Roze (2021) and Stetsenko and Roze (2022), a direct cost of recombination (parameter *c*, representing a direct fitness cost per crossover) is introduced in order to stabilize the dynamics of average map length, as indirect selection on recombination may become weaker than drift when map length is high.

## RESULTS

We first present the results from our three-locus model, and then compare multilocus extrapolations of these results with individual-based simulation results. The QLE approach consists in deriving recurrence equations describing the changes in genetic associations over one generation, and use these recursions to compute equilibrium expressions for these associations in terms of the model’s parameters and allele frequencies. As in classical deterministic models for the evolution of recombination (e.g., Barton, 1995; Lenormand, 2003), selection for recombination is driven by the linkage disequilibrium between selected loci *D*_*ab*_ (averaged over maternally and paternally inherited chromosomes), which generates associations *D*_*mab*_, *D*_*ma*_ and *D*_*mb*_ between the modifier and selected loci.

### Three-locus model

The linkage disequilibrium between alleles *a* and *b* on the maternally inherited chromosomes of females of the next generation 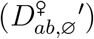 is given by:

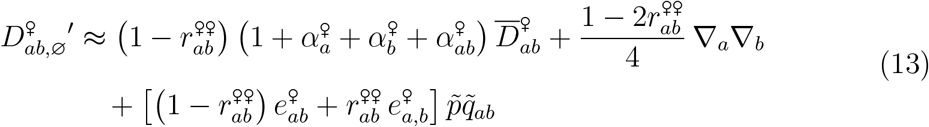

where 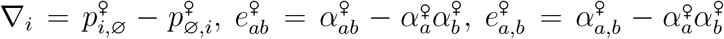 (corresponding to cis and trans epistasis measured as deviations from multiplicative selection) and 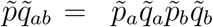 (with 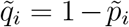). Under the fitness matrix given by Table 3, 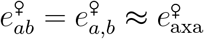 so that the last term of equation 13 simplifies to 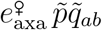. Although 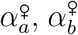 and 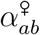 are assumed to be small, these coefficients are included in the first term of equation 13 in order to obtain more accurate approximations when recombination rates are of order *ϵ* (e.g., Roze, 2014; Stetsenko and Roze, 2022). In this tight linkage regime, equation 13 is valid to the first order in the frequencies of deleterious alleles (i.e., terms in 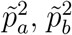 are neglected). Furthermore, the term 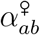 (assumed to be of order *ϵ*^2^, while 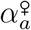 and 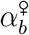 are of order *ϵ*) is included in order to improve the approximations when epistasis 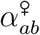 is of the same order of magnitude as the strength of effective directional selection 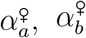. The linkage disequilibrium between *a* and *b* in males of the next generation 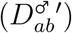 is given by the same expression, replacing 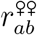 by 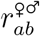, while the linkage disequilibrium on the paternally inherited chromosomes of females of the next generation is given by:

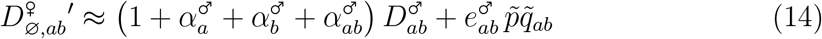

with 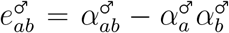. This yields the following expression for the average linkage disequilibrium in females 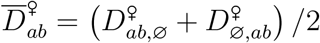 at QLE:

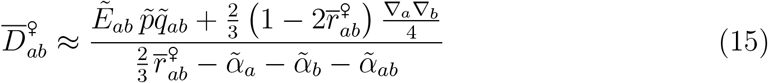

with:

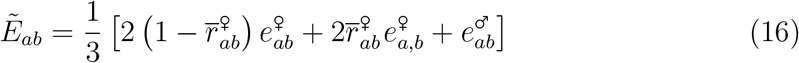

and 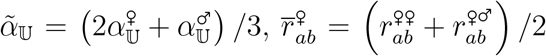. Equation 15 shows that 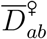 is generated by epistasis (term in 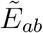, simplifying to 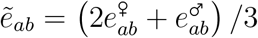 in the absence of any cis-trans difference in epistasis), and by the fact that the frequencies of *a* and *b* may differ among female and male gametes (term in ∇_*a*_∇_*b*_). It also shows that 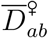 in haplodiploid species take the same form as 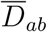 in diploid species (compare equation 15 with equation A2 in Supplementary File S1), the effect of recombination being reduced by a factor 2*/*3, since recombination does not occur in males. The effect of differences in frequencies between female and male gametes (term in ∇_*a*_∇_*b*_) is also reduced by a factor 2*/*3: this is due to the fact that male chromosomes and maternally inherited chromosomes of females are both derived from female gametes, reducing the overall effect of differences in allele frequencies between female and male gametes on LD. A last difference is that, in the haplodiploid case, the effect of selection in females is twice as important as the effect of selection in males (in 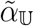 and 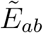 terms), while both are equally important in the diploid case.

To leading order, the changes in 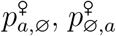 and 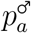 over one generation are given by:

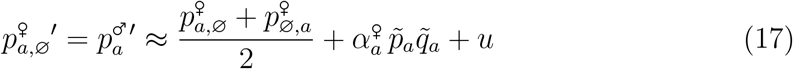

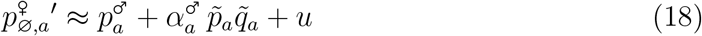

yielding, at mutation-selection balance:

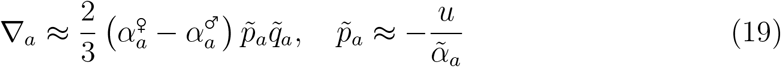

(remember that 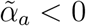) and similarly for ∇_*b*_, 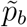. Comparing equation 19 with equation A5 in Supplementary File S1 shows that the difference in allele frequencies between female and male gametes is lower in the haplodiploid than in the diploid case (by a factor 2*/*3), for the same difference in selection coefficients between females and males. This is due to the fact that, in the haplodiploid case, all paternally inherited chromosomes were maternally inherited at the previous generation, reducing the effect of differential selection between sexes on ∇_*a*_ and ∇_*b*_.

The association between alleles *m, a* and *b* on maternally inherited chromosomes of females of the next generation is given by:

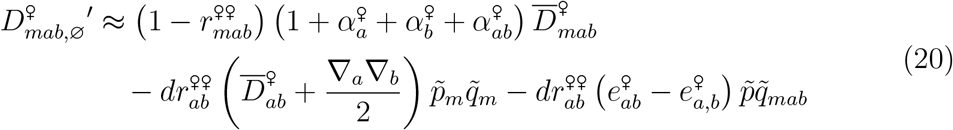

with 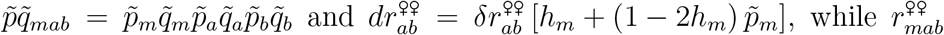 is the probability that at least one recombination event occurs between the three loci during meioses leading to daughters, given by 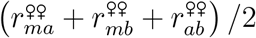 for any ordering of the three loci along the chromosome. Again, the coefficients 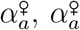 and 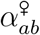 are included in the first term of equation 20 in order to obtain more accurate approximations in the case of tightly linked loci. With the fitness matrix given by Table 3, 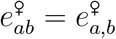 so that the last term of equation 20 vanishes. 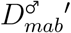 is given by the same expression, replacing 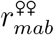 by 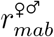 and 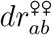 by 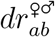, while:

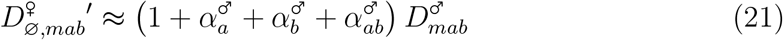

since recombination does not occur in males. From this, one obtains at QLE:

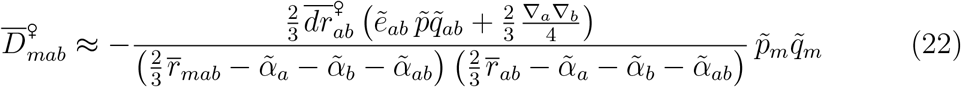

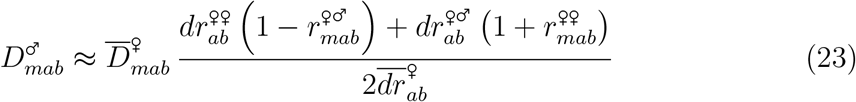

with 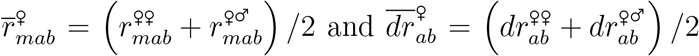. Equation 23 shows that 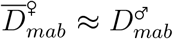 when recombination rates are identical in meioses leading to daughters and sons 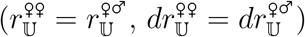. As for 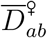 above, 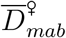 in haplodiploids takes the same form as 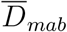 in diploids, at least in the absence of any difference between cis and trans epistasis (compare equation 22 with equation A7 in Supplementary File S1). However, recombination rates and modifier effects are multiplied by 2*/*3 (since recombination does not occur in males), while the effect of differences in allele frequencies in female and male gametes (term in ∇_*a*_∇_*b*_) is also multiplied by 2*/*3.

Finally, the association between alleles *m* and *a* on maternally inherited chromosomes of females of the next generation is given by:

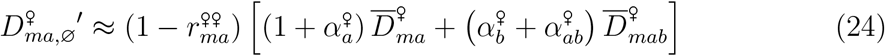

while 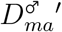 is given by the same expression after replacing 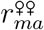 by 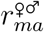, and:

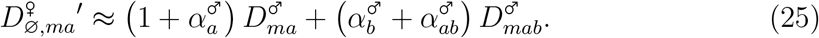

This yields, at QLE:

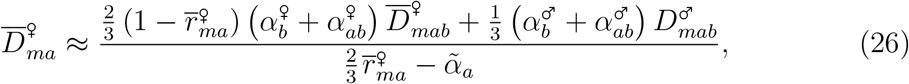

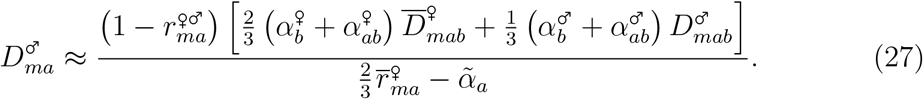

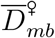 and 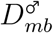 are given by the same expressions, switching *a* and *b* indices. Comparison with equation A9 in Supplementary File S1 shows that, when recombination rates are identical in meioses leading to daughters and sons (so that 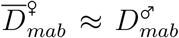 as shown above), 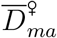 takes a similar form as 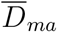 in the diploid model, recombination rates being again multiplied by 2*/*3, and selection among females having twice more effect as selection among males. Another difference is that 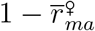 only multiplies 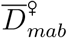 in equation 26 (and not 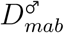, since recombination does not occur in males). However, under random mating, indirect selection on the modifier should mostly stem from selected loci that are closely linked to the modifier (e.g., Otto and Barton, 1997; Roze, 2021), in which case 1 − *r*_*ma*_ ≈ 1 and this difference between the haplodiploid and diploid cases should stay minor. Furthermore, equations 26 and 27 show that 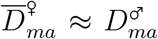 when recombination rates are identical in meioses leading to daughters and sons, at least when the recombination rate between loci *m* and *a* is sufficiently small.

The change in 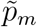 over one generation is obtained from equations 10, 22, 23, 26 and 27. When recombination rates are identical in meioses leading to daughters and sons, and to leading order in *ϵ*, we have:

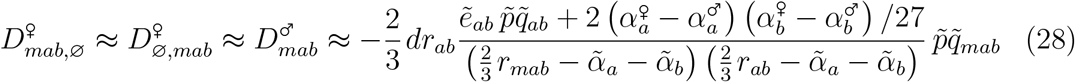

simply denoted *D*_*mab*_ hereafter, and:

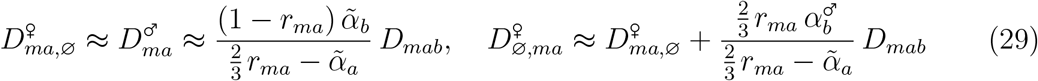

(and similarly for 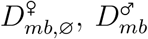 and 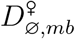, switching *a* and *b* indices). From equation 10, this yields:

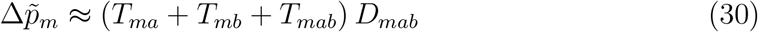

with:

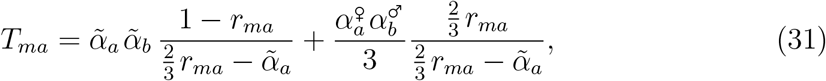

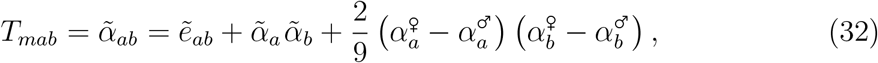

*T*_*mb*_ taking the same form as *T*_*ma*_, again switching *a* and *b* indices. When recombination rates between the modifier and selected loci are small (so that terms in *r*_*ma*_ that appear in the numerators of equation 31 may be neglected, and similarly for *T*_*mb*_), 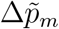 takes the same form as Δ*p*_*m*_ in a diploid model (see equations A10 – A12 in Supplementary File S1), recombination rates and the modifier effect being multiplied by 2/3, and selection coefficients in females and males being weighted by 2*/*3 and 1*/*3, respectively (instead of 1*/*2 and 1*/*2 as in the diploid case). When the effective strength of selection and epistasis at loci *a* and *b* is the same in both sexes (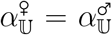 so that 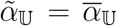), the only difference between the diploid and haplodiploid cases (again in the case where recombination rates between the modifier and selected loci are small) is that recombination rates and the modifier effect are multiplied by the 2*/*3 in the haplodiploid case. The fact that the modifier effect is multiplied by 2*/*3 reduces the magnitude of the change in frequency of *m*, while the fact that recombination rates are multiplied by 2*/*3 (in the denominators of equations 28 and 31) increases it — by increasing the strength of genetic associations. Overall, the second effect predominates, so that, when the effect of selection against deleterious alleles is identical in both sexes, indirect selection for recombination should be stronger in the haplodiploid than in the diploid case.

When selection only occurs in females 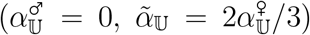, equation 30 simplifies to:

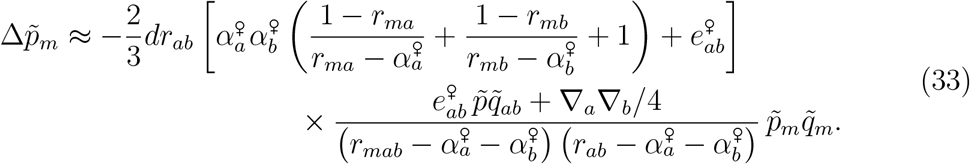

Equation 33 takes the same form as in a diploid population with equal selection in both sexes, except that the modifier effect is multiplied by 2*/*3 (since the modifier has no effect in males), and that the difference in allele frequencies between female and male gametes tends to generate positive LD (term in ∇_*a*_∇_*b*_) that disfavors recombination. Note also that under this scenario, the equilibrium frequency of deleterious alleles in the haplodiploid population would be 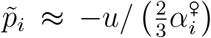, which is higher than in the case of a diploid population with equal selection in males and females (where 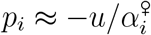), thereby increasing the magnitude of indirect selection for recombination. This difference vanishes if we assume that mutation only occurs in females of the haplodiploid population (so that males undergo no selection, no mutation and no recombination).

In the case where selection is stronger in haploid males than in diploid individuals (for example, due to fact that mutations are fully expressed in haploids), 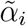 in haplodiploids should be higher in absolute value than 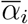 in diploids. When 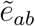 is sufficiently negative so that *D*_*mab*_ (given by equation 28) has the same sign as *dr*_*ab*_, the term (*T*_*ma*_ + *T*_*mb*_) *D*_*mab*_ of equation 30 favors recombination (effect of the increased variance in fitness among offspring), while the term *T*_*mab*_ *D*_*mab*_ disfavors it (effect of the decreased mean fitness of offspring). Because *D*_*mab*_ is proportional to 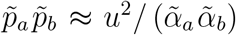 (see equation 19), the increase in absolute value of 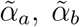 caused by stronger selection in haploid males decreases (*T*_*ma*_ + *T*_*mb*_) *D*_*mab*_ (due to the terms in 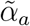 and 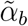 in the denominators of equations 28, 31 and the equivalent expression for *T*_*mb*_), thus decreasing the advantage of recombination. This is due to the fact that increasing the strength of selection against deleterious alleles reduces the magnitude of genetic associations, due to the selective elimination of those alleles. Assuming that 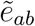 stays unchanged, increasing 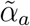 and 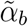 should also decrease the magnitude of the term *T*_*mab*_ *D*_*mab*_ disfavoring recombination (due to the decrease in 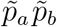, and to the terms in 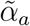 and 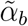 in the denominator of equations 28). In regimes where indirect selection mostly stems from selected loci that are tightly linked to the modifier locus (*r*_*ma*_, *r*_*mb*_ small), however, the term in *T*_*mab*_ should stay relatively small compared to the term in *T*_*ma*_ + *T*_*mb*_ (see Supplementary File S3), so that the main effect of increasing selection against deleterious alleles in haploid males should be to decrease the advantage of recombination stemming from increased variance in fitness among offspring. One can also note that the effect of epistasis between deleterious alleles may be stronger in haploid males than in diploid individuals (again due to the fact that mutations are fully expressed), so that 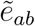 in haplodiploids may be stronger in absolute value than 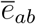 in diploids. Focusing again on the case where indirect selection mostly stems from the term *T*_*ma*_ + *T*_*mb*_ (i.e., when *r*_*ma*_, *r*_*mb*_ are small), this will increase the advantage of recombination (due to the increased absolute value of 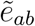 in equation 28). However, in the case of loosely linked loci and/or when 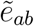 is strong relative to 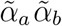 (so that the term in *T*_*mab*_ may play a more important role), an increased effect of epistasis in haploid males may disfavor recombination due to the increased recombination load (represented by the term in *T*_*mab*_).

Equation 22 shows that, to leading order in *E*, increasing recombination during meioses leading to daughters has the same effect on 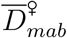 as increasing recombination during meioses leading to sons (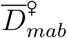 only depends on the average effect of the modifier on both types of meioses,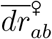). Indeed, the effect of recombination on genetic combinations in male offspring (through unfertilized ovules) is transmitted to females of the next generation, since males transmit their genotype to their daughters without recombination. By contrast, equation 23 shows that the coefficients of 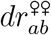 and 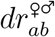 in the expression of 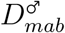 are different. In the simplest case where baseline recombination rates are identical in the two types of meiosis 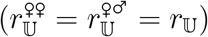, equation 23 simplifies to:

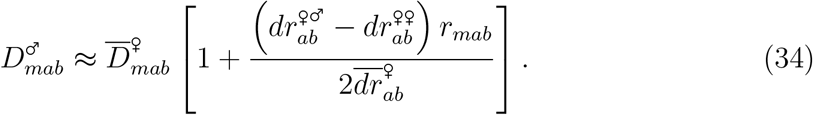

Therefore, increasing recombination during meioses leading to sons 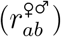 has a stronger effect on 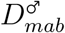 than increasing recombination during meioses leading to daughters 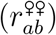. In the presence of negative epistasis, a modifier increasing 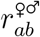 is thus more strongly disfavored than a modifier increasing 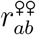 through the term 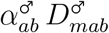 of equation 10 (representing the negative effect of decreasing the mean fitness of male offspring). However, it is also more strongly favored through the term 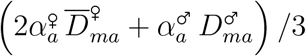 in equation 10 (and the equivalent term involving 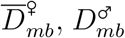) representing the positive effect stemming from increased variance in fitness among offspring. Indeed, from equations 26 – 27 and 34, this term is increased by:

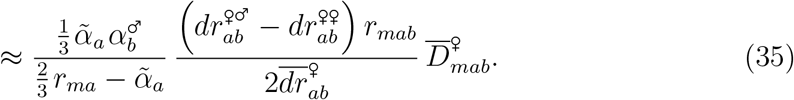

This can be understood from the fact that the benefits of recombination in terms of increased variance in fitness are greater in the case of meioses leading to male offspring (since the newly formed chromosomes are transmitted intact from male offspring to their daughters), leading to an increased efficiency of selection in one male and one female generation. This shows that the strength of selection for recombination in meioses leading to daughters and sons should generally differ, the sign of the difference potentially depending on baseline recombination rates (as the benefit of increasing the variance in fitness among offspring decreases more rapidly as recombination increases than the disadvantage of decreasing the mean fitness of offspring).

### Multilocus extrapolation and simulations

The results of our three-locus model can be extrapolated to predict the evolutionarily stable map length *R*_ES_ of a chromosome along which deleterious mutations occur at a rate *U* per generation (e.g., Roze, 2021; Stetsenko and Roze, 2022; Supplementary File S3). We first assume that the chromosome map length is the same in meioses leading to daughters and sons (controlled by a single modifier locus, and simply denoted *R* in the following), and will later consider the case where map length may differ between these two types of meioses. As in the simulations, we introduce a small direct fitness cost *c* per crossover, so that the fitness of each female is multiplied by *e*^−*cR*^ where *R* is the chromosome map length coded by its modifier locus. The change in weighted frequency of a modifier allele *m* changing map length by a small amount *δR* is given by:

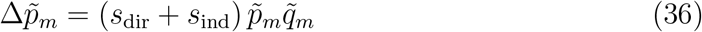

where *s*_dir_ and *s*_ind_ measure the effects of direct and indirect selection on allele *m*. Assuming that the effect of *m* is additive (*i*.*e*., the map length of *Mm* heterozygotes is *R* + *δR/*2), we have to the first order in *δR*:

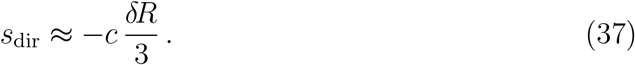

Comparing with equation A14 in Supplementary File S1 shows that direct selection is reduced compared with the diploid case (by a factor 2*/*3), due to the fact that the cost of recombination is only expressed in females. The overall effect of indirect selection *s*_ind_ is obtained by integrating the result from the three-locus model (equation 30) over all possible positions of loci *a* and *b* along the chromosome. For this, the recombination rate *r*_*ij*_ between loci *i* and *j* is expressed in terms of the genetic distance *x*_*ij*_ between these loci using Haldane’s mapping function 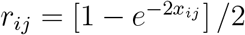 (Haldane, 1919). One obtains from this that the effect of allele *m* on *r*_*ab*_ (*dr*_*ab*_ in equation 28) is given by 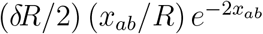 to the first order in *δR*. As shown in the *Mathematica* notebook available as Supplementary Material (https://doi.org/10.5281/zenodo.20594388), equation 30 can then be integrated numerically over all positions of selected loci along the chromosome, after replacing 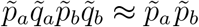 by *n*^2^, where *n* is the average number of deleterious mutations per chromosome. From equations 11 – 12 and 19, it is given by 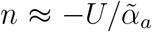, where 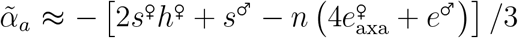— this does not take into account the difference between females and males in the mean number of mutations per chromosome, as its effect should stay negligible as long as selection against deleterious alleles is sufficiently weak. This yields:

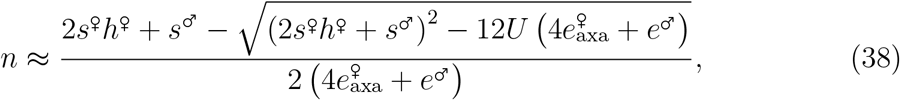

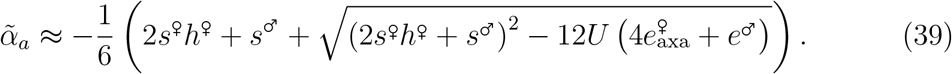

A similar extrapolation can be performed in the diploid model, as explained in Supplementary File S1. Figure 1 compares the evolutionarily stable (ES) chromosome map length *R*_ES_ predicted by our analytical model with the average map length at equilibrium measured in multilocus simulations. The analytical prediction is obtained by evaluating the overall strength of indirect selection *s*_ind_ for a range of values of *R*, and computing numerically the value of *R* for which *s*_dir_ + *s*_ind_ = 0 (as shown in the Supplementary Material). In the diploid case shown in Figure 1, selection is identical in females and males 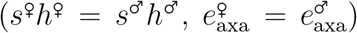, and the strength of selection *s*^♀^ = *s*^♂^ is adjusted as 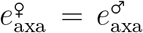 varies, in order to maintain a constant effective strength of selection against deleterious alleles 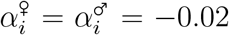 (where *i* stands for any selected locus). For *U* = 0.2 and *h*^♀^ = *h*^♂^ = 0.25 and from equations 11 and A15 in Supplementary File S1, this yields *s*^♀^ = *s*^♂^ = 0 when 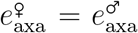 take their minimal value of −0.001 (selection is entirely driven by epistasis), while *s*^♀^ = *s*^♂^ = 0.08 when 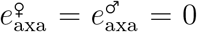 (no epistasis), *s*^♀^ and *s*^♂^ increasing linearly between these two extremes. Figure 1 shows that the ES map length increases as epistasis becomes more negative, the analytical prediction matching the simulation results except when 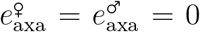. Indeed, our deterministic model predicts that indirect selection for recombination should vanish in the absence of epistasis (yielding *R*_ES_ = 0 due to the cost of recombination), while the Hill-Robertson effect caused by finite population size generates selection for recombination in the simulations (e.g., Keightley and Otto, 2006; Roze, 2021).

**Figure 1.**
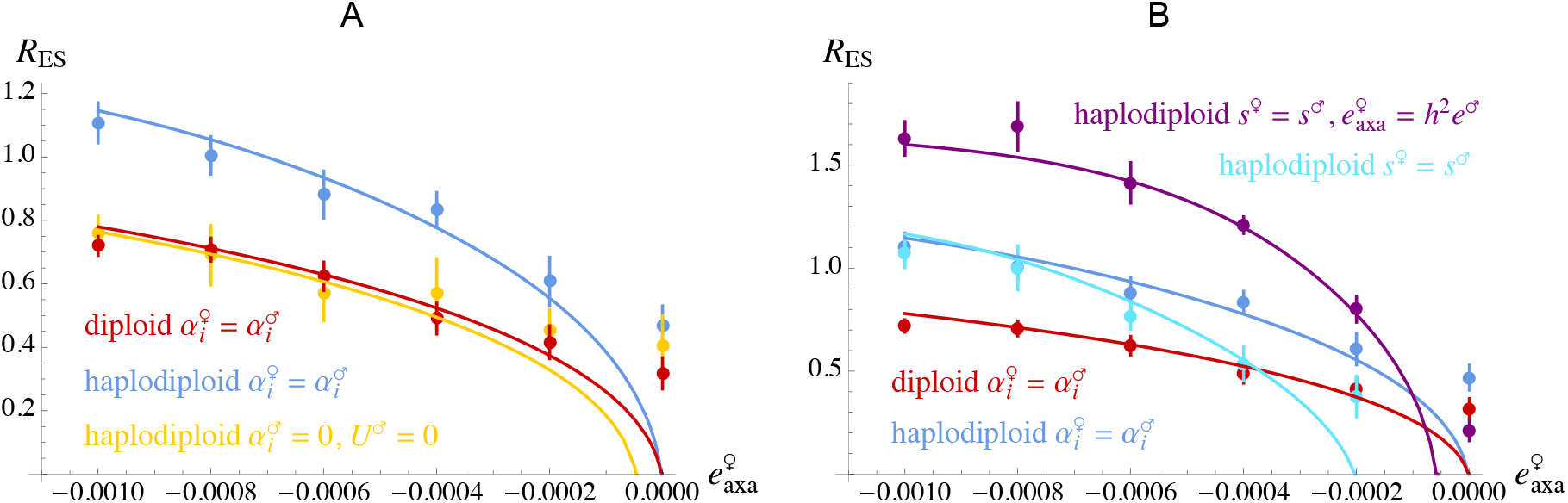
Evolutionarily stable chromosome map length *R*_ES_ (in Morgans) as a function of epistasis between pairs of deleterious alleles in females, 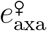. Curves: analytical predictions obtained by extrapolating our three-locus models to the case of a linear chromosome along which deleterious mutations occur at a rate *U* per generation, and where map length is controlled by a single modifier locus located at the mid-point of the chromosome (see *Mathematica* notebook available as Supplementary Material for derivations). Dots: equilibrium map length measured in multilocus individual-based simulations (a single simulation is performed for each parameter set; error bars are obtained by dividing the simulation results into batches of 5 *×* 10^5^ generations and computing the variance in average map length over batches). Parameter values: *U* = 0.2, *h*^♀^ = 0.25, *h*^♂^ = 0.25 (in diploids), *c* = 0.001, *N* = 10^4^ (in the simulations). In the”diploid 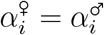” and”haplodiploid 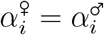” cases, *s*^♂^ and *s*^♀^ are varied as epistasis changes in order to maintain 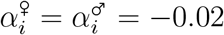. In diploids, *s*^♀^ = *s*^♂^ = 0, 0.016, 0.032, 0.048, 0.064 and 0.08 when 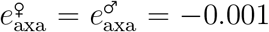, −0.0008, −0.0006, −0.0004, −0.0002 and 0, respectively. In haplodiploids, *s*^♀^, *s*^♂^ = (0, 0.01), (0.016, 0.012), (0.032, 0.014), (0.048, 0.016), (0.064, 0.018) and (0.08, 0.02) for the same values of 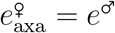. In A, *s*^♀^ stays equal to the same values in the “haplodiploid 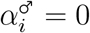, *U*^♂^ = 0 case”, while *s*^♂^ = *e*^♂^ = 0 and mutation does not occur in males. In B, *s*^♀^ = *s*^♂^ = 0, 0.016, 0.032, 0.048, 0.064 and 0.08 in the “haplodiploid *s*^♀^ = *s*^♂^” case (with 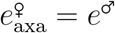), while *s*^♀^ = *s*^♂^ stay the same in the “haplodiploid *s*^♀^ = *s*^♂^, 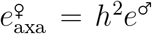” case, with *e*^♂^ = −0.016, −0.0128, −0.0096, −0.0064, −0.0032 and 0 as 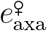 increases from −0.001 to 0. In both cases, 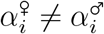, while 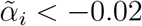.

As shown above, the results from our three-locus model indicate that in the absence of selection and mutation in haploid males, the strength of selection for recombination in haplodiploids is equivalent to its value in a diploid population where selection is identical in both sexes (and the same as in haplodiploid females) multiplied by 2*/*3 (equation in Supplementary File S1, equation 33) — it is however slightly decreased compared to the diploid case, due to the positive LD between deleterious alleles caused by the difference in selection between sexes (term in ∇_*a*_∇_*b*_ in equation 33). Because the effect of the cost of recombination is also reduced by a factor 2*/*3 in haplodiploids (compare equation A14 in Supplementary File S1 and equation 37), one predicts that the ES map length should be equivalent in both situations, but slightly lower in the haplodiploid case due to the term in ∇_*a*_∇_*b*_. As shown by Figure 1A, simulations confirm that the equilibrium chromosome map length is indeed roughly equivalent in both cases (compare “diploid 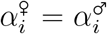” with “haplodiploid 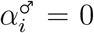, *U*^♂^ = 0” in Figure 1A). With mutation in haploid males, and when the effective strength of selection against deleterious alleles is the same in both sexes (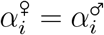, set to −0.02 in the case of Figure 1), selection for recombination is stronger in the haplodiploid than in the diploid case, however, leading to a higher map length at equilibrium (compare “diploid 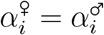” with “haplodiploid 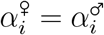” in Figure 1A). As discussed above, this results from the fact that genetic associations are stronger in the haplodiploid case (since recombination only occurs in females), increasing the effect of indirect selection. In Supplementary File S3, we show that our model predicts a 1.5 fold increase of the ES map length of haplodiploids compared with diploids due to female-limited recombination (and assuming that all else is equal between haplodiploids and diploids). This prediction is verified in Figure 1A.

In the haplodiploid case with 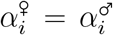 shown in Figure 1, epistasis between deleterious alleles is assumed to be the same in females and males 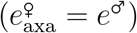. Because males are haploid, *s*^♀^ and *s*^♂^ must therefore differ in order to maintain 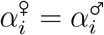: while *s*^♀^ increases linearly from 0 to 0.08 as 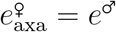 increases from −0.001 to 0 (as in the diploid case), *s*^♂^ increases linearly from 0.01 to 0.02. As shown by Figure 1B, setting *s*^♂^ equal to *s*^♀^ (also increasing from 0 to 0.08) decreases selection for recombination over most of the range of epistasis (compare “haplodiploid 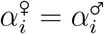” with”haplodiploid *s*^♀^ = *s*^♂^” in Figure 1B). In this case, the effect of selection against deleterious alleles is stronger in males than in females due to male haploidy 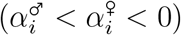 and the overall effect of selection is increased 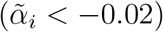, except when 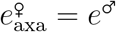 is close to −0.001 (indeed, *s*^♀^ = *s*^♂^ = 0 when 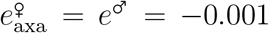, and selection against deleterious alleles is stronger in females due to fact that epistatic interactions are on average twice more numerous, see equations 11–12). A stronger effect of selection against deleterious alleles in males (due to male haploidy) thus tends to decrease selection for recombination. As shown by our three-locus model, this is due to the fact that increasing the effective strength of selection against deleterious alleles decreases the overall effect of genetic associations through the terms in *α*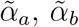 in the denominators of *T*_*ma*_, *T*_*mb*_ and *T*_*mab*_ in equation 30 (as the elimination of deleterious alleles tends to reduce the magnitude of genetic associations). Similarly, increasing the strength of selection against deleterious alleles decreases selection for recombination caused by the Hill-Robertson effect, when *Ns* is sufficiently large (Roze, 2021). Figure S1 confirms that the lower selection for recombination observed when *s*^♀^ = *s*^♂^ is caused by the increased net strength of selection against deleterious alleles (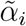 becoming more negative): indeed, setting 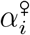 equal to 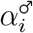, and equal to the value of 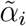 obtained in Figure 1B when *s*^♀^ = *s*^♂^ leads to a similar value of the equilibrium map length in the simulations.

While male haploidy may increase the strength of selection against deleterious alleles, it may also increase the effect of epistasis among those alleles, since epistasis may be reduced in diploids when partially recessive mutations are masked in the heterozygous state. Charlesworth et al. (1991) proposed a model of selection against deleterious alleles in which epistasis between pairs of mutations depends on their dominance coefficient when at least one of the mutations is heterozygous: in this model, epistasis between two heterozygous mutations is set to *h*^2^*e*, where *e* measures epistasis between two homozygous mutations. Under this scenario, assuming that epistasis between pairs of mutations in a haploid male is the same as epistasis among homozygous mutations in a diploid female yields 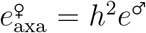 (so that epistasis is stronger in males than in females when *h <* 1). Figure 1B shows that for the same values of 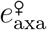, the ES chromosome map length is substantially larger when 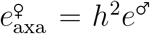 than when 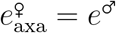, due to the stronger epistasis in males (compare “haplodiploid *s*^♀^ = *s*^♂^” with “haplodiploid *s*^♀^ = *s*^♂^, 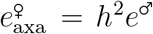” in Figure 1B). As shown by Figure S1, this is caused by the increased net effect of epistasis 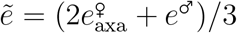: indeed, setting 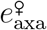 equal to *e*^♂^, and equal to the value of 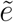 obtained in Figure 1B when 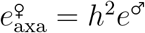 leads to the same increase in the equilibrium map length.

### Dimorphism in recombination

Our two-modifier simulation programs were used to explore the evolution of recombination dimorphism. As described in the Methods, in the diploid two-modifier program, one modifier locus controls the chromosome map length *R*^♀^ in female meioses, while the other modifier controls map length *R*^♂^ in male meioses. In the haplodiploid two-modifier program, one modifier locus controls map length *R*^♀♀^ in meioses giving rise to fertilized ovules (generating female offspring), while the other modifier controls map length *R*^♀♂^ in meioses giving rise to parthenogenetic ovules (generating male offspring). Results are shown in Figure 2. As expected, in the diploid case the average map length is the same in females and males when selection against deleterious alleles is identical in both sexes (Figure 2A), as there is no asymmetry between the sexes in this case. In haplodiploids, when the net effects of selection and epistasis are the same in both sexes, a higher map length evolves in meioses giving rise to parthenogenetic ovules than in meioses giving rise to fertilized ovules, the difference being mostly visible when 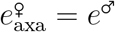 is sufficiently negative (Figure 2B). As shown in Figure 2D, this difference becomes much more important when epistasis between deleterious alleles is stronger in males (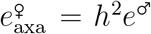, as obtained using the model of Charlesworth et al., 1991 of epistasis among deleterious alleles in diploids, assuming that epistasis in males is equivalent to epistasis between homozygous mutations in females). The curves in Figures 2B and 2D show that extrapolating our three-locus model accurately predicts this difference in evolutionarily stable map length between the two types of meioses. For this, the ES value of *R*^♀♀^ is computed for a range of values of *R*^♀♂^ (assuming that *R*^♀♂^ is fixed), while the ES value of *R*^♀♂^ is computed for a range of values of *R*^♀♀^, leading to two curves at the intersection of which lies the predicted ES value of the pair *R*^♀♀^, *R*^♀♂^ — see *Mathematica* notebook available as Supplementary Material for derivations, and Figure S2 for an example. As discussed above, the higher ES value of *R*^♀♂^ (compared with *R*^♀♀^) comes from the fact that increasing recombination in meioses leading to male offspring yields stronger benefits (in terms of increased variance in fitness among offspring) than in meioses leading to female offspring, since male genotypes are transmitted intact to their daughters.

**Figure 2.**
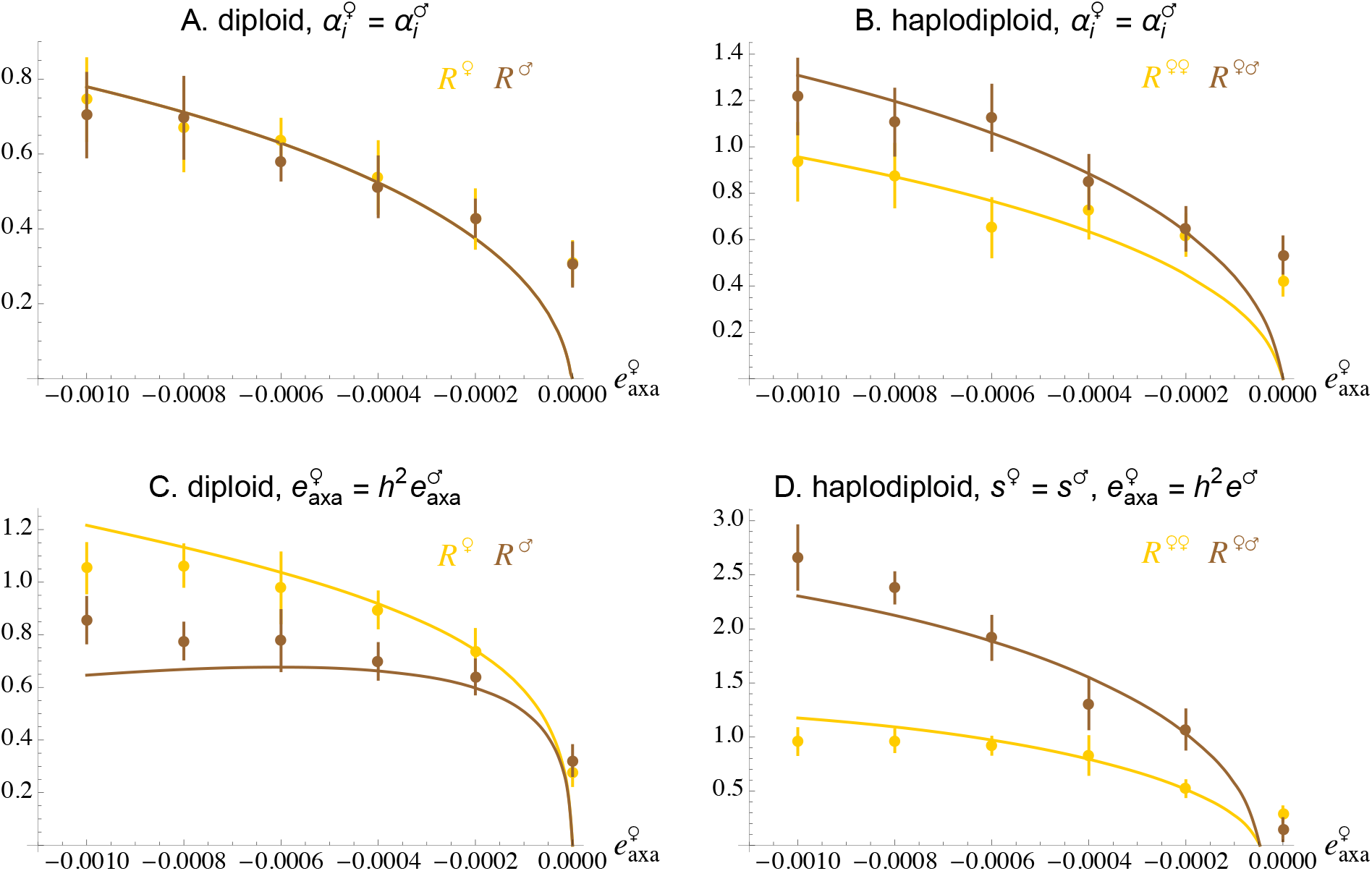
Evolutionarily stable chromosome map length (in Morgans) as a function of epistasis between pairs of deleterious alleles in females, 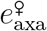, when the female and male map length (*R*^♀^ and *R*^♂^) are coded by two different modifier loci in the diploid case (A, C), while map length in meioses leading to female and male offspring (*R*^♀♀^ and *R*^♀♂^) are coded by two different loci in the haplodiploid case (B, D). Curves correspond to analytical predictions for the ES values of *R*^♀^, *R*^♂^ (diploid model) and *R*^♀♀^, *R*^♀♂^ (haplodiploid model), obtained by extrapolating our three-locus models to the case of a linear chromosome along which deleterious mutations occur at a rate *U* per generation (see *Mathematica* notebook available as Supplementary Material for derivations). Dots: equilibrium map length measured in multilocus individual-based simulations (each dot corresponds to an average over 10 replicate simulations). In A, selection is identical in both sexes (same parameter values as in Figure 1). In C, epistasis is stronger in males, with 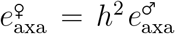. In B, the net effects of selection and epistasis between deleterious alleles are the same in both sexes (same parameter values as in Figure 1A). In D, the effect of deleterious alleles in males is the same as in homozygous females (*s*^♀^ = *s*^♂^, increasing the net effect of selection in males) while epistasis is stronger in males, with 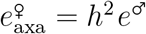.

Interestingly, Figure 2C shows that in diploids, the female map length is higher than the male map length (on average) when negative epistasis is stronger in males. This result seems to contradict the predictions of Lenormand (2003), stating that selection acting during the diploid phase of the life cycle may lead to the evolution of a sex dimorphism in recombination only if the difference between cis and trans epistasis differs between the sexes, or if the fitness effect of alleles depends on whether they are maternally or paternally inherited (none of these phenomena is present in our simulations). In Supplementary File S1 (section “sex dimorphism in recombination”), we show that the difference in indirect selection for recombination in females and males stems from the fact that the average linkage disequilibrium between deleterious alleles 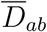 (which is the same in both sexes at the start of a generation) becomes lower in magnitude in the sex in which selection against deleterious alleles is strongest (due to the selective elimination of these alleles) at the time when recombination occurs, thereby reducing the strength of indirect selection for recombination in that sex. This effect is not taken into account in equation A6 of Supplementary File S1 (or in Lenormand, 2003), as the change in 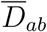 caused by selection introduces a term of higher order in *E* (and thus negligible) compared with the other terms. However, including this term leads to the prediction that selection for recombination should be stronger in females than in males for the parameter values used in Figure 2C, and extrapolating the result to the case of a whole chromosome correctly predicts the difference in map length between females and males (curves in Figure 2C, see *Mathematica* notebook available as Supplementary Material for derivation).

Figure S3 (A and D) shows that the female map length also evolves towards higher values than the male map length when sexual selection generates a form of truncation selection among males (generating negative epistasis on male fitness). In haplodiploids, however, Figure S3 (B and E) show the map length in meioses leading to sons may evolve to higher or lower values than the map length in meioses leading to daughters, depending on the intrinsic cost of recombination *c* (which determines the range of values over which map length equilibrates). When *c* = 0.01, *R*^♀♂^ evolves towards higher values than *R*^♀♀^ (as in Figures 2B and 2D), while the pattern is opposite when *c* = 0.001. This may be due to the fact that the relative importance of the disadvantageous effect of recombination (in terms of reduced mean fitness of male offspring) is stronger at higher values of map length, leading to a stronger disadvantage of modifier alleles increasing *R*^♀♂^ (compared with modifier alleles increasing *R*^♀♀^). At lower map lengths, indirect selection in mostly driven by the beneficial effect of recombination in terms of increased variance in fitness among offspring, and increasing *R*^♀♂^ leads to stronger benefits than increasing *R*^♀♀^. These simulation results could not be compared with analytical predictions, however, as computing the *α*_U, V_ selection coefficients under truncation selection is difficult. Finally, Figure S3 (C and F) show that, as in our model of pairwise epistasis among mutations, higher map lengths evolve in the haplodiploid than in the diploid case in the presence of truncation selection among males.

## DISCUSSION

Our analytical model outlines several effects of haplodiploidy on selection for recombination caused by negative epistasis among deleterious mutations. Comparing the results with those obtained under a diploid model (assuming equal recombination in both sexes), we showed that the change in frequency of a recombination modifier allele takes a similar form, the main differences being that (i) indirect selection depends on average coefficients of selection and epistasis between the sexes in the diploid case 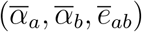 and on weighted coefficients in the haplodiploid case 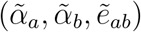, where the effect of selection among females is twice as important as selection among males; (ii) recombination rates and the modifier effect are multiplied by 2*/*3 in the haplodiploid case. Effect (i) stems from the fact that alleles spend on average twice more time in females than in males under haplodiploidy, and effect (ii) from the fact that recombination only occurs in females. When selection occurs in both sexes, the absence of recombination in males increases the strength of indirect selection on recombination by increasing the magnitude of genetic associations. However, our analysis and simulation results show that this effect alone should increase chromosome map length by a factor 3*/*2 in haplodiploids compared to diploids, which is thus not sufficient to explain the very high recombination rates observed in eusocial Hymenoptera (furthermore, this effect should also occur in non-eusocial haplodiploids, as discussed below). Another consequence of male haploidy is that it may increase the effect of selection and epistasis among males (due to the fact that mutations are fully expressed), so that 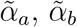 and 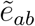 in haplodiploids may be higher in magnitude than 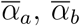 and 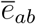 in diploids. Stronger directional selection against deleterious alleles (measured by coefficients 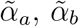) enhances the benefits of increasing the variance in fitness among offspring, but this effect is exactly compensated by the fact that it also reduces the frequency of deleterious alleles at mutation-selection balance, thereby reducing polymorphism at selected loci (*p*_*a*_*q*_*a*_*p*_*b*_*q*_*b*_) and the benefits of recombination. Selection against deleterious alleles also tends to erode genetic associations, so that overall, increasing the magnitude of coefficients 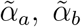 reduces the advantage of recombination under negative epistasis. However, male haploidy may also increase the overall effect of epistasis among mutations (if epistasis is stronger in haploid males, again due to the fact that mutations are fully expressed), increasing the magnitude of 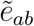. As we have seen, this selects more strongly for increased recombination when epistasis is negative. Together, the effects of female-limited recombination and stronger negative epistasis in haploid males may favor substantially higher recombination rates in haplodiploids than in diploids, as shown in Figure 1B.

Female-limited recombination and male haploidy characterize all Hymenoptera, however (not only eusocial species), while solitary Hymenoptera do not exhibit particularly high recombination rates (Niehuis et al., 2010; Jones et al., 2019; DeLory et al., 2024). Can our model explain the very high recombination rates of eusocial species? While we do not explicitly model eusociality, our results can be used to assess the plausibility of the hypothesis proposed by Everitt et al. (2025), stipulating that strong sexual selection among the males of haplodiploid eusocial species may have favored increased recombination rates. In honey bee, for example, more than 100 males may compete to fertilize a single reproductive female (e.g., Baer, 2005), and DeLory et al. (2024) showed that the recombination rate of eusocial Hymenoptera is positively correlated with the number of male mates per queen, that may be used as a proxy for the intensity of sexual selection. Furthermore, the large spermatheca of eusocial queens may provide strong opportunities for sperm competition, that may be seen as a form of sexual selection. Interestingly, termite colonies are typically derived by a single mated pair (thus reducing the opportunity for sexual selection), and the queens do not display high recombination rates (Everitt et al., 2025). However, termites belong to a different insect order from eusocial Hymenoptera and represent a single independent origin of eusociality, suggesting that their recombination rate may be constrained for other reasons.

Our results show that the effect of sexual selection on the evolution of recombination depends on the genetic architecture of fitness variation among males. If fitness variation is caused by the constant input of deleterious mutations (as in the present model), stronger selection against deleterious alleles caused by an extra effect of these alleles on male mating success (leading to an increased magnitude of coefficients 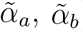) tends to decrease selection for recombination. However, sexual selection could potentially increase the magnitude of negative epistasis among mutations, which are unmasked in male haploids, thereby increasing the magnitude of 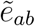 and the advantage of recombination. This would occur, for example, in a situation where only the males with relatively low numbers of deleterious alleles manage to reproduce, leading to a form of truncation selection among males. Indeed, our simulations including truncation selection on males yielded similar qualitative results as our simulations with pairwise epistasis (Figure S3). To our knowledge, the idea that sexual selection may be a source of negative epistasis among mutations has not been explored much theoretically and empirically. In haplodiploids, stronger negative epistasis caused by sexual selection may be further reinforced by male haploidy, and the combination of both effects could potentially lead to substantially elevated recombination rates.

Different results may be obtained under alternative genetic architectures of male mating success. For example, in scenarios where male fitness would depend on sweeping beneficial alleles or on alleles with sex-antagonistic effects maintained at a polymorphic equilibrium, increasing the strength of selection at these loci (due to stronger sexual selection) may increase the magnitude of directional selection coefficients 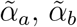 without necessarily decreasing polymorphism (*p*_*a*_*q*_*a*_*p*_*b*_*q*_*b*_), so that stronger directional selection on loci involved in male mating success may favor higher recombination rates. This could be explored further by incorporating mutations with beneficial effects on male fitness in our simulations, or considering models in which male mating success depends on a quantitative phenotypic trait whose optimal value may change over time. Such scenarios may also generate stronger overall selection for recombination (and higher evolutionarily stable chromosome map lengths) than in the mutation-selection balance scenario considered in the present article, since the variance in fitness among individuals typically stays rather low at mutation-selection balance (thereby limiting the possible strength of indirect selection at recombination modifier loci).

In our model, the advantage of recombination results from the negative LD among mutations generated by negative epistasis. However, empirical studies investigating the average sign of epistasis among deleterious mutations have yielded inconsistent results, with epistasis being reported as either positive or negative on average (Rice, 2002; Kouyos et al., 2007). Furthermore, these studies found that epistasis is typically highly variable among pairs of mutations, which is expected to disfavor recombination (Otto and Feldman, 1997). By contrast, epistasis among beneficial mutations has been more consistently found to be negative, on average (e.g., Schoustra et al., 2016; Wei and Zhang, 2019; Johnson et al., 2023). To our knowledge, the sign of epistasis among loci affecting mating success has not yet been investigated, despite its potential importance. As mentioned in the introduction, several other hypotheses have been proposed to explain the high recombination rates observed in eusocial Hymenoptera, and warrant further theoretical and empirical investigation. These include stronger benefits of recombination in facilitating the escape from parasitism (e.g., Schmid-Hempel, 2000; Sirviö et al., 2006), reducing opportunities for conflicts within colonies (Sherman, 1979; Templeton, 1979), or mitigating stronger Hill-Robertson interference resulting from the low effective population size of eusocial species (Romiguier et al., 2014; Imrit et al., 2020; Weyna and Romiguier, 2021) — indeed, higher evolutionarily stable chromosome map lengths are predicted in species with lower *N*_e_, as long as *N*_e_ is not too small (Roze, 2021). The low effective size of eusocial species also raises the possibility that high map lengths evolved due to the effect of drift at loci controlling the recombination rate, particularly if selection acting on map length is weak. A systematic increase in map length would occur through drift in the presence of a mutational bias towards higher recombination rates (that is, when mutations are more likely to increase rather than decrease map length, e.g., Zhang and Hill, 2008; Gervais and Roze, 2017; Devi et al., 2023).

Our results also show that in haplodiploids, selection for recombination differs in meioses leading to parthenogenetic ovules (generating male offspring) and in meioses leading to fertilized ovules (generating female offspring): indirect selection is stronger in the first type of meiosis, as the new haplotypes formed by recombination remain intact during one male and one female generation. Given that most linkage maps of Hymenoptera are obtained by genotyping male offspring (as it avoids the problem of phasing genotypes, since males are haploid), one may thus wonder if recombination rates may differ in meioses leading to female offspring. This may not seem very likely given the biology of oogenesis in insects, however. Indeed, while the exact timing of the different steps of meiosis has not been described yet in Hymenoptera (to our knowledge), in *Drosophila* meiosis is completed during the passage of oocytes through the oviduct, before fertilization (e.g., Avilés-Pagán and Orr-Weaver, 2018; Von Stetina and Orr-Weaver, 2011), and it may thus be difficult to adjust recombination rates according to the fate of oocytes. One can also note that some of the estimates of high recombination rates in honey bees have been obtained from patterns of linkage disequilibrium within natural populations (Wallberg et al., 2015; Jones et al., 2019), which are affected by recombination events occurring in both types of meiosis.

Finally, we have seen that in diploid species, sex differences in recombination rates (heterochiasmy) may evolve when the strength of selection differs between diploid females and males, selection for recombination being weaker in the sex in which selection against deleterious alleles is stronger. This is due to the fact that LD between selected loci is lower in magnitude in the sex in which deleterious alleles have the strongest effect, at the time where recombination occurs (as LD is lowered by selection against deleterious alleles). This effect is typically rather weak, however (leading to only moderate differences in recombination rates between the sexes, for the parameter values used in this article), and was neglected in the previous analysis by Lenormand (2003), which focused on stronger possible sources of heterochiasmy (such as sex differences in selection in the haploid phase, see also Lenormand and Duteil, 2005). In Figure 2C, selection against deleterious alleles in males is rather strong, and mainly driven by epistasis interactions with other deleterious alleles, which may not seem very plausible — for example, for 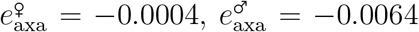 so that the direct and epistatic components of selection against deleterious alleles in males are −*s*^♂^*h*^♂^ ≈ −0.012 and 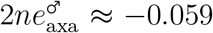. Nevertheless, this mechanism may be worth exploring further under different genetic architectures of fitness variation.

## Data availability

C++ simulation codes and *Mathematica* notebooks presenting mathematical derivations are available at https://doi.org/10.5281/zenodo.20594388.

## Acknowledgements

We thank Matt Webster, Sally Otto and two anonymous reviewers for helpful discussions and comments, and the bioinformatics and computing services at Roscoff Biological Station (Abims platform) for computing time.

