## Supplementary Figures for "Differential selection between sexes and the evolution of recombination in haplodiploids"

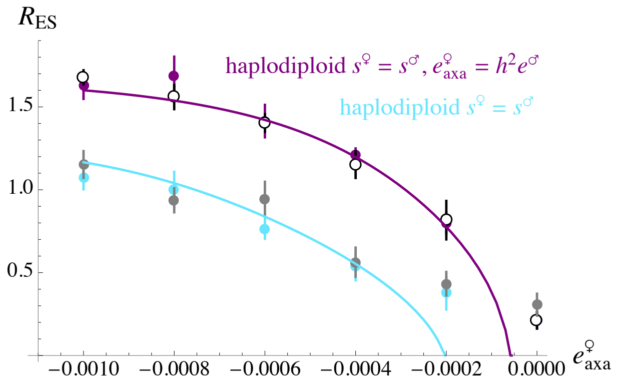


**Figure S1.** Evolutionarily stable chromosome map length $R_{ES}$ (in Morgans) in haplodiploids, as a function of epistasis between pairs of deleterious alleles in females, $e^♀_{axa}$. Coloured curves and dots for the "haplodiploid $s^♀ = s^♂$" and "haplodiploid $s^♀ = s^♂$, $e^♀_{axa} = h^2 e^♂$" cases are the same as in Figure 1B. Grey dots correspond to simulation results obtained for $e^♂ = e^♀_{axa}$, the values of $s^♀$ and $s^♂$ being set so that $\alpha_i^♀ = \alpha_i^♂ = \tilde{\alpha}_i$ while the value of $\tilde{\alpha}_i$ is the same as in the simulations shown in the "haplodiploid $s^♀ = s^♂$" case, in which $\alpha_i^♀ \neq \alpha_i^♂$. For this, and using equations 12–13 and 56, $\left(s^♀, s^♂\right) = (-0.015, 0.0073)$, $(0.022, 0.013)$, $(0.057, 0.019)$, $(0.092, 0.026)$, $(0.126, 0.033)$ and $(0.16, 0.04)$ for $e^♀_{axa} = e^♂ = -0.001, -0.0008, -0.0006, -0.0004, -0.0002$ and 0 (respectively), yielding $\tilde{\alpha}_i \approx -0.018, -0.021, -0.024, -0.028, -0.034$ and $-0.04$, as in the $s^♀ = s^♂$ case. White dots correspond to simulation results obtained for $e^♂ = e^♀_{axa} = \tilde{e}$, the values of $s^♀$, $s^♂$ and $\tilde{e}$ being set so that $\alpha_i^♀$, $\alpha_i^♂$ and $\tilde{e}$ are the same as in the simulations shown in the "haplodiploid $s^♀ = s^♂$, $e^♀_{axa} = h^2 e^♂$" case, in which $e^♀_{axa} \neq e^♂$. For this, $\left(s^♀, s^♂\right) = (-0.219, 0.055)$, $(-0.157, 0.059)$, $(-0.096, 0.064)$, $(-0.036, 0.069)$, $(0.023, 0.074)$ and $(0.08, 0.08)$ while $e^♀_{axa} = e^♂ = \tilde{e} = -0.006, -0.0048, -0.0036, -0.0024, -0.0012$ and 0, yielding $\tilde{\alpha}_i \approx -0.0365, -0.0369, -0.0374, -0.0380, -0.0389$ and $-0.04$, as in the "haplodiploid $s^♀ = s^♂$, $e^♀_{axa} = h^2 e^♂$" case.

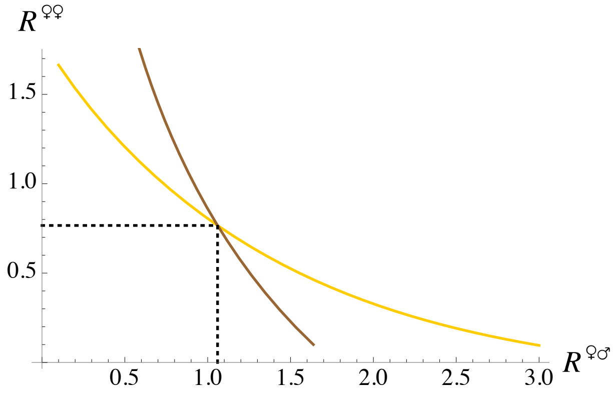


**Figure S2.** Analytical predictions of evolutionarily stable (ES) map lengths in meioses leading to fertilized and unfertilized ovules ($R^{♀♀}$ and $R^{♀♂}$, respectively) in haplodiploids, obtained by extrapolating our three-locus model to the case of a whole chromosome. Parameter values are as in Figure 2B with $e^{♀}_{\text{axa}} = -0.0006$. Yellow curve: ES value of $R^{♀♀}$ (y-axis) as a function of $R^{♀♂}$ (x-axis); brown: ES value of $R^{♀♂}$ (x-axis) as a function of $R^{♀♀}$ (y-axis). Curves are obtained by interpolation over 30 points (see *Mathematica* notebook available on Zenodo, https://doi.org/10.5281/zenodo.20594388 for more details). $R^{♀♀}$ is predicted to increase below the yellow curve and to decrease above the curve, while $R^{♀♂}$ is predicted to increase on the left of the brown curve and to decrease on the right of the curve. As a consequence, the predicted ES value of the $\left(R^{♀♀}, R^{♀♂}\right)$ pair lies at the intersection between the two curves.

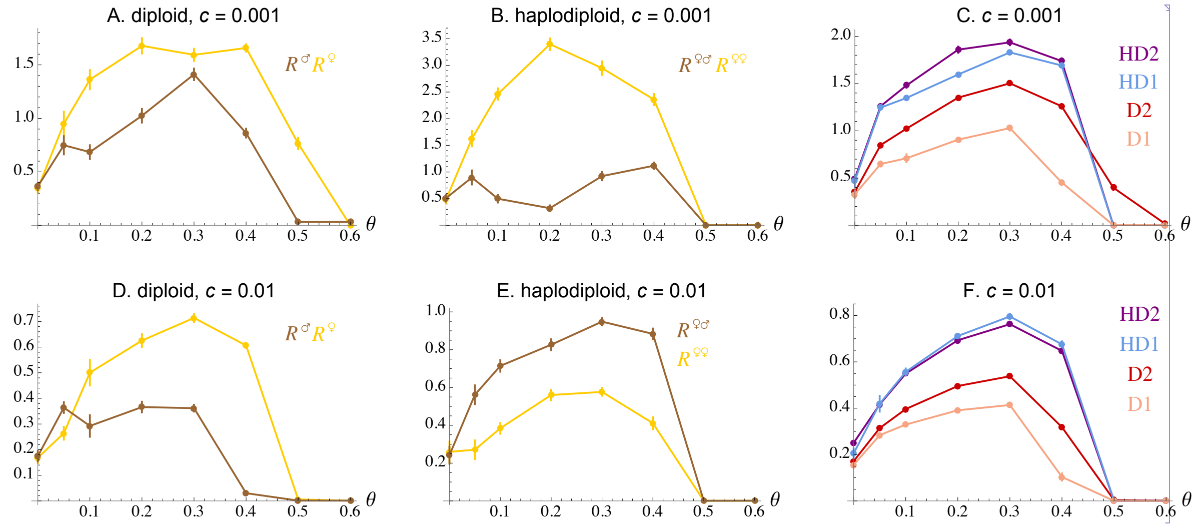


**Figure S3.** Average map length at equilibrium observed in multilocus simulations, in the presence of truncation selection among males, whose intensity is measured by $\theta$ (x-axes): the $N^{♂}\theta$ males with the lowest fitness (obtained by assuming multiplicative effects of deleterious alleles at different loci) are excluded from reproduction (while selection is multiplicative in females). The direct fitness cost of recombination is $c = 0.001$ in top figures (A, B, C) and $c = 0.01$ in bottom figures (D, E, F). A, D: diploid population, two-modifier model (the female and male map length $R^{♀}$ and $R^{♂}$ evolve independently). B, E: haplodiploid population, two-modifier model (the map length in meioses leading to female and male offspring, $R^{♀♀}$ and $R^{♀♂}$, evolve independently). C, F: average map length over all types of meioses in the two-modifier simulations shown on the left figures (D2, HD2), and average map length at equilibrium in one-modifier simulations (D1, HD1) where map length is the same in both sexes (diploid model) or in meioses leading to fertilized and unfertilized ovules (haplodiploid case). Parameter values are $N = 10^4$, $s^{♀} = s^{♂} = 0.02$, $h^{♀} = h^{♂} = 0.25$, $U = 0.2$. The colored lines simply connect simulation results and do not correspond to analytical predictions.
