## Supplementary File S1 for "Differential selection between sexes and the evolution of recombination in haplodiploids"

FILE S1: QLE ANALYSIS OF THE DIPLOID MODEL

We describe here the analysis of the diploid model, assuming that selection and recombination may differ between females and males. Our results are equivalent to those obtained by Lenormand (2003), in the absence of selection occurring in the haploid phase of the life cycle and of differential fitness effects of maternally and paternally inherited alleles (these effects are considered by Lenormand, 2003). However, some higher-order terms are retained here in order to obtain more accurate approximations in the case of tightly linked loci, that we can then integrate along a linear chromosome (following Roze, 2021; Stetsenko and Roze, 2022). Because we assume that all loci are autosomal, at the start of a generation allele frequencies and genetic associations are necessarily the same in females in males. In particular, $p^{♀}_{i,\varnothing} = p^{♂}_{i,\varnothing}$ and $p^{♀}_{\varnothing,i} = p^{♂}_{\varnothing,i}$, so that the frequency of allele $i$ in the whole population is given by $p_i = \left(p^{♀}_{i,\varnothing} + p^{♀}_{\varnothing,i}\right)/2 = \left(p^{♂}_{i,\varnothing} + p^{♂}_{\varnothing,i}\right)/2$. Furthermore, $D^{♀}_{\mathbb{U},\mathbb{V}} = D^{♂}_{\mathbb{U},\mathbb{V}}$ (that we will simply denote $D_{\mathbb{U},\mathbb{V}}$) at the start of a generation. Throughout the following, $\overline{D}_{\mathbb{U}}$ is defined as $\overline{D}_{\mathbb{U}} = \left(D_{\mathbb{U},\varnothing} + D_{\varnothing,\mathbb{U}}\right)/2$. As shown by equation 9 in the main text, the change in frequency of the modifier can be expressed in terms of associations $\overline{D}_{ma}$, $\overline{D}_{mb}$ and $\overline{D}_{mab}$, that are computed in the following section.

**Three-locus model.** The linkage disequilibrium between alleles $a$ and $b$ on the maternally inherited chromosomes of the next generation ($D'_{ab,\varnothing}$) is given by:

$$\begin{aligned} D'_{ab,\varnothing} \approx \left(1 - r^{♀}_{ab}\right)\left(1 + \alpha^{♀}_{a} + \alpha^{♀}_{b} + \alpha^{♀}_{ab}\right)\overline{D}_{ab} + \frac{1 - 2r^{♀}_{ab}}{4}\nabla_a\nabla_b \\ + \left[\left(1 - r^{♀}_{ab}\right)e^{♀}_{ab} + r^{♀}_{ab}\,e^{♀}_{a,b}\right]pq_{ab} \end{aligned} \tag{A1}$$

where $\nabla_i = p_{i,\varnothing} - p_{\varnothing,i}$, $e^{♀}_{ab} = \alpha^{♀}_{ab} - \alpha^{♀}_{a}\alpha^{♀}_{b}$, $e^{♀}_{a,b} = \alpha^{♀}_{a,b} - \alpha^{♀}_{a}\alpha^{♀}_{b}$ (corresponding to cis and trans epistasis measured as deviations from multiplicativity) and $pq_{ab} = p_a q_a p_b q_b$ (with $q_i = 1 - p_i$). Under the fitness matrix given by Table 3, $e^{♀}_{ab} = e^{♀}_{a,b} \approx e^{♀}_{\mathrm{axa}}$ when $p_a$ and $p_b$ are small, so that the last term of equation A1 simplifies to $e^{♀}_{\mathrm{axa}}\,pq_{ab}$. The linkage disequilibrium between alleles $a$ and $b$ on the paternally inherited chromosomes of the next generation ($D'_{\varnothing,ab}$) is given by the same expression, replacing female by male symbols. From this, one obtains the following expression for $\overline{D}_{ab}$ at QLE:

$$\overline{D}_{ab} \approx \frac{E_{ab}\,pq_{ab} + \left(1 - 2\overline{r}_{ab}\right)\frac{\nabla_a\nabla_b}{4}}{\overline{r}_{ab} - \overline{\alpha}_a - \overline{\alpha}_b - \overline{\alpha}_{ab}} \tag{A2}$$

where $\overline{\alpha}_{\mathbb{U}} = \left(\alpha^{♀}_{\mathbb{U}} + \alpha^{♂}_{\mathbb{U}}\right)/2$, $\overline{r}_{ab} = \left(r^{♀}_{ab} + r^{♂}_{ab}\right)/2$, and

$$E_{ab} = \frac{1}{2}\left[\left(1 - r^{♀}_{ab}\right) e^{♀}_{ab} + r^{♀}_{ab}\, e^{♀}_{a,b}\right] + \frac{1}{2}\left[\left(1 - r^{♂}_{ab}\right) e^{♂}_{ab} + r^{♂}_{ab}\, e^{♂}_{a,b}\right] \tag{A3}$$

simplifying to $\overline{e}_{ab} = \left(e^{♀}_{ab} + e^{♂}_{ab}\right)/2$ when cis and trans epistasis are identical (as in the fitness matrix given in Table 3). Equation A2 shows that $\overline{D}_{ab}$ is generated by epistasis (term in $E_{ab}$), and by the fact that the frequencies of $a$ and $b$ may differ among female and male gametes (term in $\nabla_a \nabla_b$). To leading order in $\epsilon$ and $u$, the changes in $p_{a,\varnothing}$ and $p_{\varnothing,a}$ over one generation are given by:

$$p'_{a,\varnothing} \approx p_a + \alpha^{♀}_{a}\, p_a q_a + u, \quad p'_{\varnothing,a} \approx p_a + \alpha^{♂}_{a}\, p_a q_a + u \tag{A4}$$

(and similarly for $p_{b,\varnothing}$, $p_{\varnothing,b}$), so that:

$$\nabla_a \approx \left(\alpha^{♀}_{a} - \alpha^{♂}_{a}\right) p_a q_a, \quad p_a \approx -u/\overline{\alpha}_a \tag{A5}$$

at mutation-selection balance (and similarly for $\nabla_b$, $p_b$). When $\overline{r}_{ab} \gg \epsilon$, equation A2 is equivalent to the result given in Lenormand (2003) (his equation 19, extended to the case where recombination rates differ between sexes as explained on the top right of p. 817, and replacing 4 by 2 on the second line of his equation 20, which may be due to a typo in this equation). Note that the term $\overline{\epsilon}^{\mathrm{D}}_{jk}$ defined by equation 21 of Lenormand (2003) is equivalent to $\overline{e}_{ab} + \left(\alpha^{♀}_{a} - \alpha^{♂}_{a}\right)\left(\alpha^{♀}_{b} - \alpha^{♂}_{b}\right)/4$.

An expression for the association between alleles $m$, $a$ and $b$ on maternally inherited chromosomes of the next generation is given by:

$$\begin{aligned} D'_{mab,\varnothing} \approx{} & \left(1 - r^{♀}_{mab}\right)\left(1 + \alpha^{♀}_{a} + \alpha^{♀}_{b} + \alpha^{♀}_{ab}\right) \overline{D}_{mab} \\ & - dr^{♀}_{ab}\left(\overline{D}_{ab} + \frac{\nabla_a \nabla_b}{2}\right) p_m q_m - dr^{♀}_{ab}\left(e^{♀}_{ab} - e^{♀}_{a,b}\right) pq_{mab} \end{aligned} \tag{A6}$$

with $pq_{mab} = p_m q_m p_a q_a p_b q_b$ and $dr^{♀}_{ab} = \delta r^{♀}_{ab}\left[h_m + (1 - 2h_m)\, p_m\right]$, while $r^{♀}_{mab}$ is the probability that at least one recombination event occurs between the three loci during a female meiosis, given by $\left(r^{♀}_{ma} + r^{♀}_{mb} + r^{♀}_{ab}\right)/2$ for any ordering of the three loci along the chromosome. With the fitness matrix given by Table 3, $e^{♀}_{ab} = e^{♀}_{a,b}$ so that the last term of equation A6 vanishes. $D'_{\varnothing,mab}$ is given by the same expression, replacing female by male symbols. This yields the following expression for $\overline{D}_{mab}$ at QLE:

$$\begin{aligned} \overline{D}_{mab} \approx -\frac{1}{\overline{r}_{mab} - \overline{\alpha}_a - \overline{\alpha}_b - \overline{\alpha}_{ab}} \Bigg[ & \overline{dr}_{ab}\, \frac{E_{ab}\, pq_{ab} + \frac{\nabla_a \nabla_b}{4}}{\overline{r}_{ab} - \overline{\alpha}_a - \overline{\alpha}_b - \overline{\alpha}_{ab}}\, p_m q_m \\ & + \left[\frac{dr^{♀}_{ab}}{2}\left(e^{♀}_{ab} - e^{♀}_{a,b}\right) + \frac{dr^{♂}_{ab}}{2}\left(e^{♂}_{ab} - e^{♂}_{a,b}\right)\right] pq_{mab} \Bigg] \end{aligned} \tag{A7}$$

where $\overline{r}_{mab} = \left(r^{♀}_{mab} + r^{♂}_{mab}\right)/2$ and $\overline{dr}_{ab} = \left(dr^{♀}_{ab} + dr^{♂}_{ab}\right)/2$. Finally, the linkage disequilibrium between alleles $m$ and $a$ on the maternally inherited chromosome of the next generation is given by:

$$D'_{ma,\varnothing} \approx \left(1 - r^{♀}_{ma}\right)\left[\left(1 + \alpha^{♀}_{a}\right)\overline{D}_{ma} + \left(\alpha^{♀}_{b} + \alpha^{♀}_{ab}\right)\overline{D}_{mab}\right] \tag{A8}$$

$D'_{\varnothing,ma}$ being again given by the same expression after replacing female by male symbols, yielding at QLE:

$$\overline{D}_{ma} \approx \frac{1}{2}\frac{\left(1 - r^{♀}_{ma}\right)\left(\alpha^{♀}_{b} + \alpha^{♀}_{ab}\right) + \left(1 - r^{♂}_{ma}\right)\left(\alpha^{♂}_{b} + \alpha^{♂}_{ab}\right)}{\overline{r}_{ma} - \overline{\alpha}_{a}}\,\overline{D}_{mab}. \tag{A9}$$

$\overline{D}_{mb}$ at QLE is given by the same expression, switching $a$ and $b$ indices.

The change in frequency of the modifier allele $m$ is obtained from equations 9 in the main text, A7 and A9. When baseline recombination rates are the same in females and males ($r^{♀}_{\mathbb{U}} = r^{♂}_{\mathbb{U}}$) and when $r_{\mathbb{U}} \gg \epsilon$, the result obtained can be shown to be equivalent to equation 25 in Lenormand (2003) in the absence of haploid selection or sex-of-origin effect. When recombination rates and modifier effects are the same in females and males ($r^{♀}_{\mathbb{U}} = r^{♂}_{\mathbb{U}}$, $\delta r^{♀}_{\mathbb{U}} = \delta r^{♂}_{\mathbb{U}}$), when cis and trans epistasis are identical, and to leading order in $\epsilon$, this simplifies to:

$$\Delta p_m \approx \left(T_{ma} + T_{mb} + T_{mab}\right)\overline{D}_{mab} \tag{A10}$$

with:

$$\overline{D}_{mab} = -dr_{ab}\,\frac{\overline{e}_{ab} + \left(\alpha^{♀}_{a} - \alpha^{♂}_{a}\right)\left(\alpha^{♀}_{b} - \alpha^{♂}_{b}\right)/4}{\left(r_{mab} - \overline{\alpha}_{a} - \overline{\alpha}_{b}\right)\left(r_{ab} - \overline{\alpha}_{a} - \overline{\alpha}_{b}\right)}\,pq_{mab}, \tag{A11}$$

$$T_{ma} = \overline{\alpha}_{a}\overline{\alpha}_{b}\,\frac{1 - r_{ma}}{r_{ma} - \overline{\alpha}_{a}}, \quad T_{mab} = \overline{e}_{ab} + \overline{\alpha}_{a}\overline{\alpha}_{b} + \frac{\left(\alpha^{♀}_{a} - \alpha^{♂}_{a}\right)\left(\alpha^{♀}_{b} - \alpha^{♂}_{b}\right)}{4}, \tag{A12}$$

$T_{mb}$ taking the same form as $T_{ma}$, switching $a$ and $b$ indices. When selection is identical in both sexes and when $r_{\mathbb{U}} \gg \epsilon$, equations A10 – A12 are equivalent to equation A1.5$f$ in Barton (1995), showing that increased recombination is favored when epistasis is negative and linkage between the modifier and selected loci is sufficiently tight (so that $T_{ma} + T_{mb} + T_{mab} > 0$). The same result holds in the presence of sex differences in selection, replacing selection coefficients by their averages between sexes, and replacing epistasis by $\overline{e}_{ab} + \left(\alpha^{♀}_{a} - \alpha^{♂}_{a}\right)\left(\alpha^{♀}_{b} - \alpha^{♂}_{b}\right)/4$. Indeed, epistasis and differences in selection between sexes both generate LD among gametes (explaining why they both appear on the numerator of equation A11), and also affect the difference in marginal fitness of coupling and repulsion gametes (coefficient $\overline{\alpha}_{ab}$, corresponding to the term $T_{mab}$). This shows that in the absence of epistasis, increased recombination can be favored when $\left(\alpha_a^{♀} - \alpha_a^{♂}\right)\left(\alpha_b^{♀} - \alpha_b^{♂}\right)$ is negative (for example, if allele $a$ is more deleterious in females while allele $b$ is more deleterious in males) and linkage is sufficiently tight.

Differences between sexes in recombination rates may evolve when the coefficients of $dr_{ab}^{♀}$ and $dr_{ab}^{♂}$ differ in the expression of the change in frequency of the modifier (meaning that modifiers affecting recombination rates in females and males are differently selected). From equation A7, this arises when $e_{ab}^{♀} - e_{a,b}^{♀} \neq e_{ab}^{♂} - e_{a,b}^{♂}$, that is, when there is a difference between cis and trans epistasis, and this difference differs between sexes. This result was obtained by Lenormand (2003), while Sardell and Kirkpatrick (2020) proposed a mechanism (involving interactions between mutations in genes with sex-specific expression and their cis regulatory sequences) that may generate such a difference. Lenormand (2003) also showed that sex differences in recombination may result from sex differences in selection in the haploid phase, or from sex-of-origin effects during diploid selection (*i.e.*, when the fitness effects of alleles depend on whether they are maternally or paternally inherited). Indeed, in both cases, the fitness consequences of breaking LD between selected loci differ between female and male meioses.

**Multilocus extrapolation.** As explained in the main text, the results of this three-locus model can be extrapolated to predict the evolutionarily stable map length $R_{\mathrm{ES}}$ of a chromosome along which deleterious mutations occur at a rate $U$ per generation. In the following, we assume a uniform density of mutations and crossovers along the chromosome (without crossover interference) and introduce a small direct fitness cost $c$ per crossover, so that the fitness of each individual is multiplied by $e^{-cR}$ where $R$ is the chromosome map length coded by its modifier locus. When map length is the same in females and males (and controlled by a single modifier locus), the change in frequency of a modifier allele $m$ changing $R$ by a small amount $\delta R$ is given by:

$$\Delta p_m = \left(s_{\mathrm{dir}} + s_{\mathrm{ind}}\right) p_m q_m \tag{A13}$$

where $s_{\mathrm{dir}}$ and $s_{\mathrm{ind}}$ measure the effects of direct and indirect selection on allele $m$. Assuming that the effect of $m$ is additive (*i.e.*, the map length of $Mm$ heterozygotes is $R + \delta R/2$), we have to the first order in $\delta R$:

$$s_{\text{dir}} \approx -c\,\frac{\delta R}{2} \tag{A14}$$

(Roze, 2021; Stetsenko and Roze, 2022), while $s_{\text{ind}}$ is obtained by integrating the result from the three-locus model (equation A10) over all possible positions of loci $a$ and $b$ along the chromosome (see main text and *Mathematica* notebook available as Supplementary Material, https://doi.org/10.5281/zenodo.20594388). For this, equation A10 is integrated numerically over all positions of selected loci along the chromosome, after replacing $p_a q_a p_b q_b \approx p_a p_b$ by $n^2$, where $n$ is the average number of deleterious mutations per chromosome. From equation 11 in the main text and equation A5, it is given by $n \approx -U/\overline{\alpha}_a$ where $\overline{\alpha}_a \approx -\overline{sh} + 2n\overline{e}$ and $\overline{sh} = \left(s^{♀}h^{♀} + s^{♂}h^{♂}\right)/2$, $\overline{e} = \left(e^{♀}_{\text{axa}} + e^{♂}_{\text{axa}}\right)/2$, yielding:

$$n \approx \frac{1}{4\overline{e}}\left(\overline{sh} - \sqrt{\overline{sh}^2 - 8U\overline{e}}\right), \quad \overline{\alpha}_a \approx -\frac{1}{2}\left(\overline{sh} + \sqrt{\overline{sh}^2 - 8U\overline{e}}\right). \tag{A15}$$

**Sex dimorphism in recombination.** The analysis of the three-locus model presented above shows that the strength of indirect selection on recombination should be the same in female and male meioses, unless the difference between cis and trans epistasis differs between sexes ($e^{♀}_{ab} - e^{♀}_{a,b} \neq e^{♂}_{ab} - e^{♂}_{a,b}$). However, cis and trans epistasis are identical in our simulations, so that $e^{♀}_{ab} - e^{♀}_{a,b} = e^{♂}_{ab} - e^{♂}_{a,b} = 0$. In this case, equation A7 shows that indirect selection only depends on $\overline{dr}_{ab} = \left(dr^{♀}_{ab} + dr^{♂}_{ab}\right)/2$, so that the strength of indirect selection on a mutation increasing recombination in females should be the same as on a mutation having the same effect in males.

Differences in the strength of indirect selection acting on mutations affecting recombination in females and males emerge when higher-order terms are included in the recursions on genetic associations, however. In particular, a more accurate version of equation A6 is given by:

$$\begin{aligned} D'_{mab,\varnothing} \approx {} & \left(1 - r^{♀}_{mab}\right)\left(1 + \alpha^{♀}_a + \alpha^{♀}_b + \alpha^{♀}_{ab}\right)\overline{D}_{mab} \\ & - dr^{♀}_{ab}\left[\left(1 + \alpha^{♀}_a + \alpha^{♀}_b + \alpha^{♀}_{ab}\right)\overline{D}_{ab} + \frac{\nabla_a \nabla_b}{2}\right] p_m q_m \end{aligned} \tag{A16}$$

(where cis and trans epistasis are assumed to be equal). The association between $m$, $a$ and $b$ on paternally inherited chromosomes ($D'_{\varnothing,mab}$) is given by the same expression, replacing female by male symbols. The term $\left(\alpha_a^{\female} + \alpha_b^{\female} + \alpha_{ab}^{\female}\right) \overline{D}_{ab}$ that appears in equation A16 was ignored in equation A6 (as in the analysis of Lenormand, 2003), as it is of higher order in $\epsilon$ than $\overline{D}_{ab}$. This term represents the fact that the linkage disequilibrium between deleterious alleles $a$ and $b$ is decreased (in absolute value) by the effect of selection against those alleles ($\alpha_a^{\female}$, $\alpha_b^{\female}$ and $\alpha_{ab}^{\female}$ are negative), thus reducing the strength of selection for recombination. Therefore, indirect selection on an allele increasing recombination in males is weaker than on an allele having the same effect in females when selection against deleterious alleles is stronger in males, due to the fact that the linkage disequilibrium between deleterious alleles is weaker in males at the time when recombination occurs.

Another potential source of asymmetry of indirect selection between the sexes stems from how direct selection on recombination is implemented in our simulation model. Indeed, we assume a direct fitness cost of crossovers, the fitnesses of females and males being multiplied by factors $e^{-cR^{\female}}$ and $e^{-cR^{\male}}$ (respectively), where $R^{\female}$ and $R^{\male}$ correspond to the map length of a focal female or male. As a result, the frequency of a modifier allele $m$ on maternally and paternally inherited chromosomes will differ if this allele has different effects on female and male map length ($\delta R^{\female}$, $\delta R^{\male}$). To leading order, this difference is given by:

$$\nabla_m = p_{m,\varnothing} - p_{\varnothing,m} \approx -\frac{c}{2}\left(\delta R^{\female} - \delta R^{\male}\right) p_m q_m. \tag{A17}$$

Similarly, we have seen that the frequency of the deleterious allele $a$ differs on maternally and paternally inherited chromosomes when selection differs between sexes, with $\nabla_a \approx \left(\alpha_a^{\female} - \alpha_a^{\male}\right) p_a q_a$ (equation A5). Differences in allele frequencies at loci $m$ and $a$ between maternally and paternally inherited haplotypes generate linkage disequilibrium when these haplotypes are combined in the same diploid individuals. From equation A2, the resulting linkage disequilibrium is approximately:

$$\overline{D}_{ma} \approx \frac{\left(1 - 2\overline{r}_{ma}\right) \nabla_m \nabla_a}{4\left(\overline{r}_{ma} - \overline{\alpha}_a\right)}. \tag{A18}$$

Therefore, when selection against allele $a$ is stronger in males ($\alpha_a^{\male} < \alpha_a^{\female} < 0$ so that $\nabla_a > 0$), this effect generates negative LD between a modifier allele increasing recombination in females (but having no effect in males) and the deleterious allele $a$, and positive LD between a modifier allele increasing recombination in males (but having no effect in females) and the deleterious allele. This mechanism is thus another source of asymmetry between modifiers acting on female vs. male recombination, increasing the strength of indirect selection on modifier alleles increasing female recombination (relative to modifier alleles increasing male recombination) when selection against deleterious alleles is stronger in males.

Similarly, one can show that the difference in frequency of $m$ between maternally and paternally inherited haplotypes ($\nabla_m$) generates a three-locus association $\overline{D}_{mab}$ when the LD between deleterious alleles also differs between those haplotypes ($D_{ab,\varnothing} - D_{\varnothing,ab} \neq 0$), which is generally the case when epistasis differs between sexes. When epistasis is negative and stronger in males, this tends to generate negative $\overline{D}_{mab}$ when allele $m$ increases recombination in females (but has no effect in males), and positive $\overline{D}_{mab}$ when $m$ increases recombination in males (but has no effect in females). However, numerical analyses indicate that this source of asymmetry is generally negligible relative to the two other sources discussed above (results not shown).

The following figure shows the results of multilocus simulations in which only two modifier alleles segregate, either at the locus controlling female map length $R^{♀}$ (orange) or at the locus controlling male map length $R^{♂}$ (brown), in the case where epistasis is negative and stronger in males (parameter values correspond to Figure 2C with $e^{♀}_{\text{axa}} = -0.0006$). The baseline map length is the same in both sexes ($R$, shown on the $x$-axis), while the mutant allele increases either female or male map length by an amount $\delta R = 0.1$. Recurrent mutation between the two modifier alleles occurs at rate $10^{-4}$, maintaining polymorphism. Dots on the figure show the covariance between the genetically encoded map length of the individual and the number of deleterious alleles in its genome (measured over all females in simulations in which the modifier locus affecting female map length is segregating, and over all males in simulations in which the modifier locus affecting male map length is segregating), divided by the variance in genetically encoded map length. This quantity can be shown to be equivalent to $\sum_i \overline{D}_{mi} / \left[(\delta R/2)\, p_m q_m\right]$, where the sum is over all selected loci. Solid curves show the analytical predictions obtained by integrating the result from the three-locus model over all possible positions of deleterious alleles $a$ and $b$ along the chromosome, taking into account the reduction in $\overline{D}_{ab}$ caused by selection in the expression of $\overline{D}_{mab}$ (equation A16) and the linkage disequilibrium caused by the term in $\nabla_m \nabla_a$ (equation A18). Dotted curves correspond to the same prediction, but neglecting the term in $\nabla_m \nabla_a$. The simulations confirm that, when negative epistasis is stronger in males, a mutant increasing recombination in females is slightly more advantaged than a mutant increasing recombination in males, due to more negative LD between the modifier and deleterious alleles. Furthermore, the difference in the strength of indirect selection acting on these two mutants is well predicted by our analytical model, when the baseline map length $R$ is sufficiently large. Finally, it shows that this difference is mostly due to the different effects of selection on $\overline{D}_{ab}$ in males and females, the term in $\nabla_m \nabla_a$ shown in equation A18 making only a minor contribution (compare solid and dotted curves).

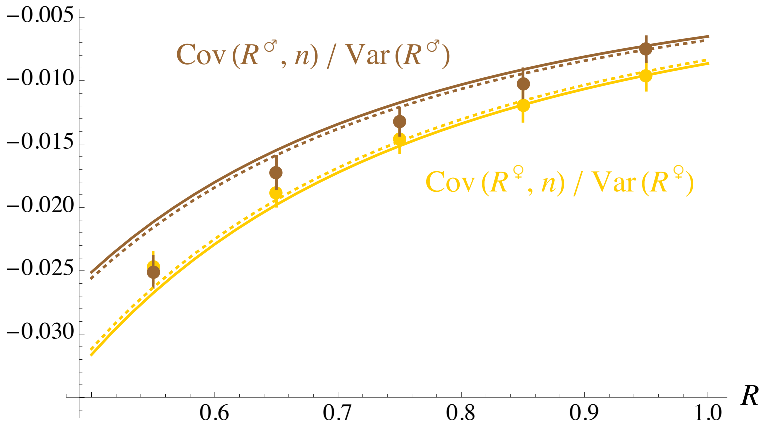


Covariance between the genetically encoded map length of individuals and the number of deleterious alleles they carry, scaled by the variance in genetically encoded map length (equivalent to $\sum_i \overline{D}_{mi} / [(\delta R/2)\, p_m q_m]$). Dots: multilocus simulations in which only two modifier alleles segregate, either at the modifier locus affecting female map length (orange) or at the modifier locus affecting male map length (brown). Each dot is an average over 60 replicate simulations, each simulation lasting $5 \times 10^6$ generations. Curves: analytical predictions (see text for more explanations). Parameter values are as in Figure 2C with $e^{♀}_{\text{axa}} = -0.0006$, $e^{♂}_{\text{axa}} = -0.0096$, and where the baseline chromosome map length of females and males in shown on the $x$-axis (the mutant modifier allele increasing map length by an amount $\delta R = 0.1$).
