## Supplementary File S2 for "Differential selection between sexes and the evolution of recombination in haplodiploids"

FILE S2: MULTILOCUS SIMULATION PROGRAMS

Our individual-based simulation programs (written in C++ and available from *Zenodo*, https://doi.org/10.5281/zenodo.20594388) represent a population consisting in $N$ individuals, each carrying either one (if haploid) or two (if diploid) copies of a linear chromosome. Selection and recombination occur as in the programs used in Roze (2021) and Stetsenko and Roze (2022): each generation, the number of new deleterious mutations per chromosome is drawn from a Poisson distribution with parameter $U$, and the position of each new mutation along the chromosome is drawn from a uniform distribution. The fitness of a diploid individual of sex $S$ (either ♀ or ♂) is computed as:

$$W^S = \left(1 - s^S h^S\right)^{n_{\mathrm{He}}} \left(1 - s^S\right)^{n_{\mathrm{Ho}}} \left(1 + e^S_{\mathrm{axa}}\right)^{n_{\mathrm{P}}} \tag{B1}$$

where $n_{\mathrm{He}}$ and $n_{\mathrm{Ho}}$ are the numbers of heterozygous and homozygous mutations in the genome of the individual, while $n_{\mathrm{P}}$ is the number of pairs of mutations at different loci, given by $n_{\mathrm{tot}}\left(n_{\mathrm{tot}} - 1\right)/2 - n_{\mathrm{Ho}}$ where $n_{\mathrm{tot}} = n_{\mathrm{He}} + 2n_{\mathrm{Ho}}$. In the haplodiploid model, the fitness of haploid males is computed as:

$$W^{♂} = \left(1 - s^{♂}\right)^n \left(1 + e^{♂}\right)^{\frac{n(n-1)}{2}} \tag{B2}$$

where $n$ is the number of mutations in the genome of the individual. To produce the individuals of the next generation, parents are drawn with probabilities corresponding to their fitness divided by the mean fitness of individuals from the same sex. Modified versions of our programs were used to consider the case where sexual selection generates a form a truncation selection among males. For this, the fitnesses of males and females were computed using equations B1 and B2 without epistasis ($e^{♀}_{\mathrm{axa}} = e^{♂}_{\mathrm{axa}} = 0$ in the diploid case, $e^{♀}_{\mathrm{axa}} = e^{♂} = 0$ in the haplodiploid case), and the $N^{♂}\theta$ males with the lowest fitness were not allowed to reproduce, where $N^{♂}$ is the number of males in the population (the remaining proportion $1-\theta$ of males contribute to the next generation in proportion to their fitness). Such truncation selection generates negative epistasis among deleterious alleles on male reproductive success (e.g., Kondrashov, 1993).

During meiosis, the number of crossovers is drawn from a Poisson distribution whose parameter depends on the values of modifier alleles present in the individual, while the position of each crossover is drawn from a uniform distribution along the chromosome. In the diploid model, each newly formed individual becomes female or male with equal probabilities, while in the haplodiploid model $N/2$ females are produced by fertilization, while $N/2$ males are produced from unfertilized ovules from selected female parents. In both the diploid and haplodiploid model, chromosome map length is controlled by either one or two modifier loci. In the one-modifier case, map length is the same in males and females (in the diploid model) or in meioses giving rise to female and male offspring (in the haplodiploid model), and the modifier locus is located at the mid-point of the chromosome. In the two-modifier case, one modifier locus controls map length in female meioses (diploid model) or in meioses giving rise to female offspring (haplodiploid model), while the other modifier controls map length in male meioses or in meioses giving rise to male offspring. In that case, the modifier loci are located at 1/3 and 2/3 of the length of the chromosome.

During the first $2 \times 10^4$ generations, map length is fixed to $R = 1$ in order to reach mutation-selection balance for deleterious alleles. Then, mutation is introduced at modifier loci (at a rate $10^{-4}$ per locus per generation). When a mutation occurs, with probability 0.95 the value of the map length coded by the modifier allele is multiplied by a number drawn from a Gaussian distribution with average 1 and variance 0.04, while with probability 0.05 a number drawn from a uniform distribution between $-1$ and 1 is added to the value coded by the allele, in order to allow for both small and large-effect mutations (map length is set to zero if the new value is negative). For simplicity, modifier alleles have additive effects on map length. As in Roze (2021) and Stetsenko and Roze (2022), a direct fitness cost of recombination was introduced in order to stabilize the dynamics of average map length (as indirect selection on recombination may become weaker than drift when map length is high). For this, the fitness of diploid individuals is multiplied by $e^{-cR}$, where $R$ is the chromosome map length of the individual (in the haplodiploid two-modifier model, $R$ is the average between $R^{♀♀}$ and $R^{♀♂}$). The parameter $c$ thus represents a direct fitness cost per crossover.
