## Supplementary File S3 for "Differential selection between sexes and the evolution of recombination in haplodiploids"

### FILE S3: EVOLUTIONARILY STABLE MAP LENGTH IN THE HAPLODIPLOID AND DIPLOID MODELS

As discussed in the main text, indirect selection on recombination in the diploid and haplodiploid cases take similar forms when the effective strength of selection against deleterious alleles (and epistasis) is identical in diploid females, haploid males and diploid males (so that $\tilde{\alpha}_{\mathbb{U}}$ in the haplodiploid model and $\overline{\alpha}_{\mathbb{U}}$ in the diploid model are identical), the only effect of haplodiploidy being that recombination rates and the modifier effect are multiplied by 2/3 in the expression for the change in frequency of the modifier allele (since recombination occurs only in females). In this file, we show that under these assumptions, and assuming a fixed direct fitness cost of recombination, the evolutionarily stable (ES) chromosome map length in haplodiploids should be approximately 1.5 times the value of the ES map length in diploids.

Assuming that recombination rates are the same in meioses leading to fertilized and unfertilized ovules and that epistasis is weak relative to selection against deleterious alleles ($\tilde{\alpha}_{ab} \ll \tilde{\alpha}_a$, $\tilde{\alpha}_b$), an approximate expression for the change in frequency of the modifier caused by indirect selection in given by equations 28 and 30–32 in the main text. Under weak selection, indirect selection mostly stems from selected loci located in the genomic vicinity of the modifier locus ($r_{ma}$, $r_{mb}$ small), so that the terms in $r_{ma}$ in the numerators of equation 31 may be neglected. Furthermore, the term $T_{mab}$ of equation 30 is negligible relative to $T_{ma}$, $T_{mb}$ when loci are tightly linked. When the effective strength of selection is the same in females and males ($\alpha_a^{♀} = \alpha_a^{♂}$, $\alpha_b^{♀} = \alpha_b^{♂}$), and using the fact that $\tilde{p}_a\tilde{q}_a \approx -u/\tilde{\alpha}_a$, $\tilde{p}_b\tilde{q}_b \approx -u/\tilde{\alpha}_b$ (equation 19), the change in frequency of the modifier caused by indirect selection simplifies to:

$$\Delta\tilde{p}_m \approx -\frac{2}{3} dr_{ab} \frac{\tilde{e}_{ab}\,\tilde{p}_m\tilde{q}_m}{\left(\frac{2}{3}r_{mab} - \tilde{\alpha}_a - \tilde{\alpha}_b\right)\left(\frac{2}{3}r_{ab} - \tilde{\alpha}_a - \tilde{\alpha}_b\right)} \left[\frac{1}{\frac{2}{3}r_{ma} - \tilde{\alpha}_a} + \frac{1}{\frac{2}{3}r_{mb} - \tilde{\alpha}_b}\right] \quad \text{(C1)}$$

with $r_{mab} = \left(r_{ma} + r_{mb} + r_{ab}\right)/2$. Extrapolating to the situation where deleterious alleles occur along a whole chromosome, this expression can be integrated over all possible positions of pairs of deleterious alleles to obtain the overall effect of indirect selection on a modifier affecting the total map length of the chromosome (here simply denoted $R$). The curves shown in the figures of the article have been obtained by expressing recombination rates $r_{ij}$ in terms of genetic distances $x_{ij}$ using Haldane's mapping function: $r_{ij} = \left(1 - e^{-2x_{ij}}\right)/2$, leading to $dr_{ab} \approx (\delta R/2)\,(x_{ab}/R)\,e^{-2x_{ab}}$ to the first order in the modifier effect on map length $\delta R$, and integrating over genetic distances. As will be illustrated below, approximating recombination rates by genetic distances ($r_{ij} \approx x_{ij}$, $dr_{ab} \approx (\delta R/2)\,(x_{ab}/R)$) yields very similar results, again due to the fact that indirect selection is mostly generated by tightly linked loci ($x_{ij}$ small). Using this approximation and taking into account the effect of the direct cost of recombination (given by equation 37 in the main text), one obtains the following expression for the change in frequency of the modifier:

$$\Delta \tilde{p}_m \approx -\frac{2}{3}\frac{\delta R}{2}\left[c + \tilde{e}_{ab}\,U^2\,(2I_1 + I_2)\right]\tilde{p}_m \tilde{q}_m \tag{C2}$$

with:

$$\begin{aligned} I_1 = \frac{1}{R^3}\int_0^{\frac{R}{2}}\int_0^{\frac{R}{2}-x_{ma}} & \frac{x_{ab}}{\left[\frac{2}{3}\left(x_{ma}+x_{ab}\right)-2\tilde{\alpha}_a\right]\left[\frac{2}{3}x_{ab}-2\tilde{\alpha}_a\right]} \\ & \times \left[\frac{1}{\frac{2}{3}x_{ma}-\tilde{\alpha}_a}+\frac{1}{\frac{2}{3}\left(x_{ma}+x_{ab}\right)-\tilde{\alpha}_a}\right] dx_{ab}\,dx_{ma} \end{aligned} \tag{C3}$$

$$I_2 = \frac{1}{R^3}\int_0^{\frac{R}{2}}\int_0^{\frac{R}{2}} \frac{x_{ma}+x_{mb}}{\left[\frac{2}{3}\left(x_{ma}+x_{mb}\right)-2\tilde{\alpha}_a\right]^2}\left[\frac{1}{\frac{2}{3}x_{ma}-\tilde{\alpha}_a}+\frac{1}{\frac{2}{3}x_{mb}-\tilde{\alpha}_a}\right] dx_{ma}\,dx_{mb} \tag{C4}$$

where $I_1$ is an integral over all pairs of loci located on the same side of the chromosome (left or right, the modifier locus being located at the mid-point of the chromosome, i.e., loci in the order $m - a - b$), while $I_2$ is an integral over all pairs of loci located on opposite sides of the modifier locus (loci in the order $a - m - b$). The integrands of $I_1$ and $I_2$ can be simplified using the change in variables $y_{ij} = -x_{ij}/\tilde{\alpha}_a$, yielding:

$$\begin{aligned} I_1 = \frac{1}{R^3}\int_0^{-\frac{R}{2\tilde{\alpha}_a}}\int_0^{-\frac{R}{2\tilde{\alpha}_a}-y_{ma}} & \frac{y_{ab}}{\left[\frac{2}{3}\left(y_{ma}+y_{ab}\right)+2\right]\left[\frac{2}{3}y_{ab}+2\right]} \\ & \times \left[\frac{1}{\frac{2}{3}y_{ma}+1}+\frac{1}{\frac{2}{3}\left(y_{ma}+y_{ab}\right)+1}\right] dy_{ab}\,dy_{ma} \end{aligned} \tag{C5}$$

$$I_2 = \frac{1}{R^3}\int_0^{-\frac{R}{2\tilde{\alpha}_a}}\int_0^{-\frac{R}{2\tilde{\alpha}_a}} \frac{y_{ma}+y_{mb}}{\left[\frac{2}{3}\left(y_{ma}+y_{mb}\right)+2\right]^2}\left[\frac{1}{\frac{2}{3}y_{ma}+1}+\frac{1}{\frac{2}{3}y_{mb}+1}\right] dy_{ma}\,dy_{mb}\,. \tag{C6}$$

The next figure shows that the overall strength of indirect selection on recombination obtained from equations C2, C5 and C6 is very close to the expression obtained by integrating equations 28 and 30–32 in the main text (including the term $T_{mab}$, and without approximating recombination rates by genetic distances).

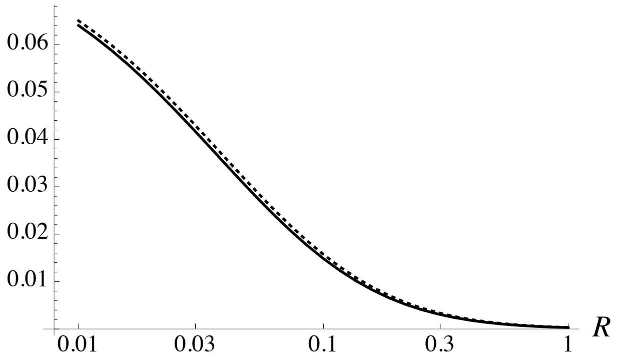


Strength of indirect selection on recombination (measured as $\Delta\tilde{p}_m / (\delta R\, \tilde{p}_m \tilde{q}_m)$ when $c = 0$) obtained from equations C2, C5 and C6 (dotted curve), and by integrating equations 28 and 30–32 in the main text (solid curve), as a function of the baseline chromosome map length $R$ (on a log scale). Parameter values are the same as in Figure 2 in the main text for the "haploidiploid $\alpha_i^{♀} = \alpha_i^{♂}$" case, with $e_{\mathrm{axa}}^{♀} = e^{♂} = -0.0006$, $\alpha_a^{♀} = \alpha_a^{♂} = -0.02$, $U = 0.2$.

Under the same hypotheses, the change in frequency of a modifier of map length in a diploid species is given by the same expression, replacing $\tilde{e}_{ab}$ by $\overline{e}_{ab}$ and $\tilde{\alpha}_a$ by $\overline{\alpha}_a$, and removing the 2/3 factor in equation C2, and the 2/3 factors in the denominators of the integrands of $I_1$ and $I_2$. In the haplodiploid case, using the change in variables $z_{ij} = (2/3)y_{ij}$ yields:

$$I_1 = \frac{1}{\left(\frac{2}{3}R\right)^3} \int_0^{-\frac{\frac{2}{3}R}{2\tilde{\alpha}_a}} \int_0^{-\frac{\frac{2}{3}R}{2\tilde{\alpha}_a} - z_{ma}} \frac{z_{ab}}{[(z_{ma} + z_{ab}) + 2]\,[z_{ab} + 2]} \times \left[\frac{1}{z_{ma} + 1} + \frac{1}{(z_{ma} + z_{ab}) + 1}\right] dz_{ab}\, dz_{ma} \tag{C7}$$

$$I_2 = \frac{1}{\left(\frac{2}{3}R\right)^3} \int_0^{-\frac{\frac{2}{3}R}{2\tilde{\alpha}_a}} \int_0^{-\frac{\frac{2}{3}R}{2\tilde{\alpha}_a}} \frac{z_{ma} + z_{mb}}{[(z_{ma} + z_{mb}) + 2]^2} \left[\frac{1}{z_{ma} + 1} + \frac{1}{z_{mb} + 1}\right] dz_{ma}\, dz_{mb}\,. \tag{C8}$$

These integrals are the same as those obtained in the diploid model, except that $R$ is changed to $(2/3)R$. The evolutionarily stable (ES) map length corresponds to the value of $R$ for which the term between brackets of equation C2 equals zero (indirect and direct selection exactly cancel each other). Therefore, $(2/3)\,R_{\mathrm{ES}}$ in the haplodiploid case should be the same as $R_{\mathrm{ES}}$ in the diploid case (where $R_{\mathrm{ES}}$ corresponds to the evolutionarily stable map length), when $\tilde{e}_{ab}$ and $\tilde{\alpha}_a$ in the haplodiploid model are the same as $\overline{e}_{ab}$ and $\overline{\alpha}_a$ in the diploid model.
